# Binding affinity landscapes constrain the evolution of broadly neutralizing anti-influenza antibodies

**DOI:** 10.1101/2021.05.25.445596

**Authors:** Angela M. Phillips, Katherine R. Lawrence, Alief Moulana, Thomas Dupic, Jeffrey Chang, Milo S. Johnson, Ivana Cvijović, Thierry Mora, Aleksandra M. Walczak, Michael M. Desai

## Abstract

Over the past two decades, several broadly neutralizing antibodies (bnAbs) that confer protection against diverse influenza strains have been isolated^1,2^. Structural and biochemical characterization of these bnAbs has provided molecular insight into how they bind distinct antigens^1^. However, our understanding of the evolutionary pathways leading to bnAbs, and thus how best to elicit them, remains limited. Here, we measure equilibrium dissociation constants of combinatorially complete mutational libraries for two naturally isolated influenza bnAbs^3–5^ (CR-9114, 16 mutations; CR-6261, 11 mutations), reconstructing all possible intermediates back to the unmutated germline sequences. We find that these two libraries exhibit strikingly different patterns of breadth: while many variants of CR-6261 display moderate affinity to diverse antigens, those of CR-9114 display appreciable affinity only in specific, nested combinations. By examining the extensive pairwise and higher-order epistasis between mutations, we find key sites with strong synergistic interactions that are highly similar across antigens for CR-6261 and different for CR-9114. Together, these features of the binding affinity landscapes strongly favor sequential acquisition of affinity to diverse antigens for CR-9114, while the acquisition of breadth to more similar antigens for CR-6261 is less constrained. These results, if generalizable to other bnAbs, may explain the molecular basis for the widespread observation that sequential exposure favors greater breadth^6–8^, and such mechanistic insight will be essential for predicting and eliciting broadly protective immune responses.

---

Vaccination harnesses the adaptive immune system, which responds to new pathogens by mutating antibody-encoding genes and selecting for variants that bind the pathogen of interest. However, influenza remains a challenging target for immunization: most antibodies elicited by vaccines provide protection against only a subset of strains, largely due to the rapid evolution of the influenza surface protein hemagglutinin (HA)^9,10^. After nearly two decades of studies, only a handful of broadly neutralizing antibodies (bnAbs) have been isolated from humans, with varying degrees of cross-protection against diverse strains^1,3,4,11^. Still, we do not fully understand many factors affecting how and when bnAbs are produced. In particular, affinity is acquired through a complex process of mutation and selection^12^, but the effects of mutations on binding affinity to diverse antigens are not well characterized.

For example, consider two well-studied influenza bnAbs that display varying levels of breadth: CR-9114 is one of the broadest anti-influenza antibodies ever found, neutralizing strains from both groups of influenza A and strains from influenza B, while CR-6261 is limited to neutralizing strains from Group 1 of influenza A^3–5,13^. Both antibodies were isolated from vaccinated donors, derive from very similar germline sequences (IGHV1–69 and IGHJ6), and bind the conserved HA stem epitope (ED Fig. 1)^3–5^. Each antibody heavy chain has many mutations (18 amino acid changes for CR-9114, 14 for CR-6261, Fig. 1a), including seven positions that are mutated in both, yet the contributions of these mutations to affinity against different antigens remain unclear^4^.

**Figure 1:**
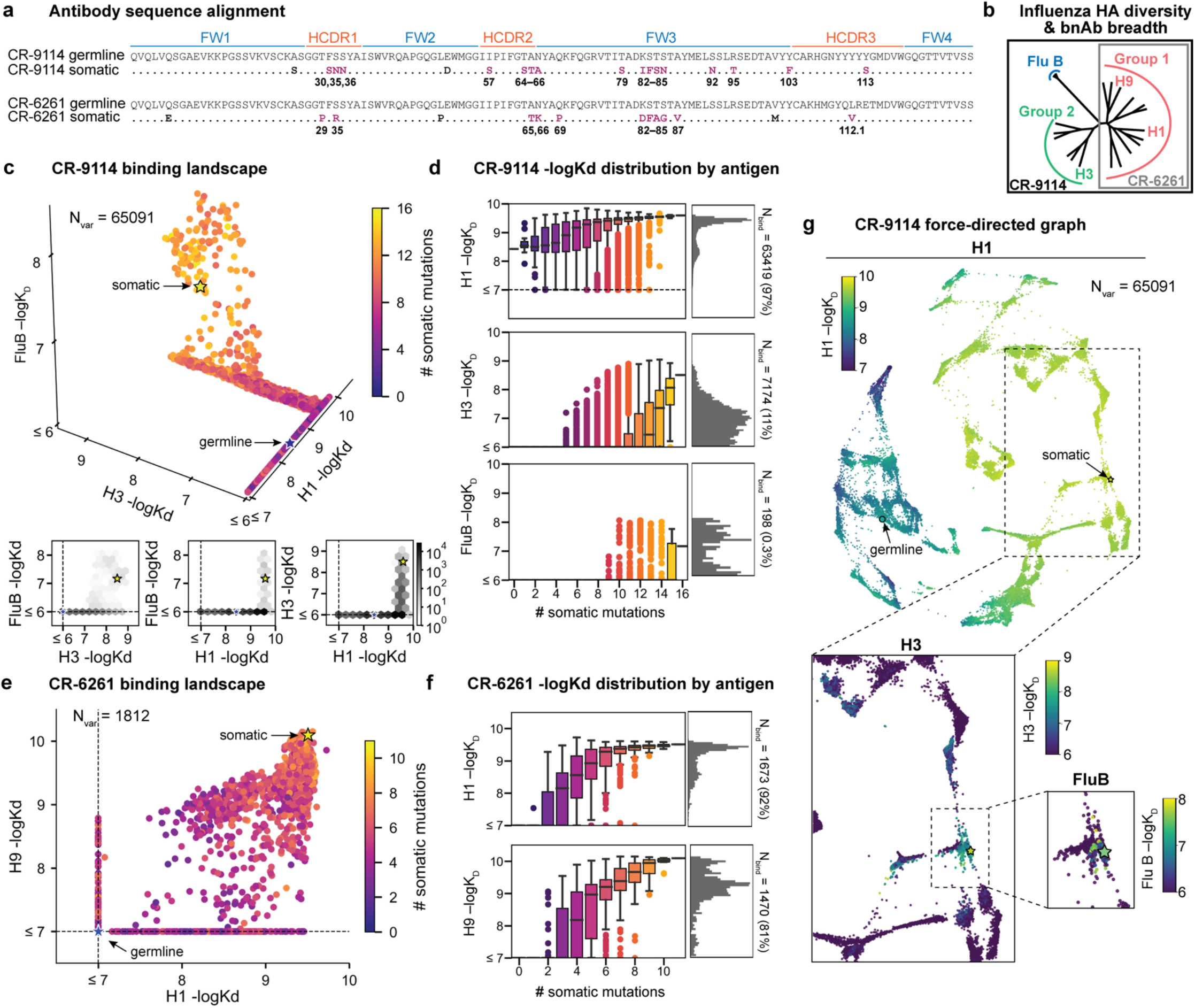
Binding landscapes. **a,** Sequence alignment comparing somatic heavy chains to reconstructed germline sequences. Mutations under study (purple, numbered) and excluded mutations (black) are indicated. **b,** Influenza hemagglutinin phylogenetic tree with selected antigens and breadth of CR-9114 (black box) and CR-6261 (gray box) indicated. **c, e,** Scatterplots of the **(c)** CR-9114 library binding affinities against three antigens, with 2D planes shown below, and **(e)** CR-6261 library binding affinities against two antigens. **d, f,** Distributions of library binding affinities for **(d)** CR-9114 and **(f)** CR-6261 for each antigen (grey histogram, right) separated by number of somatic mutations (boxplots, left). Numbers and percentages of variants with measurable binding are indicated at right. **g**, Force-directed graph of CR-9114 H1 −logK_D_. Each variant (node) is connected to its 16 single-mutation neighbors (edges not shown for clarity); edges are weighted such that variants with similar genotypes and −logK_D_ tend to cluster. Nodes are colored by binding affinity to H1 (top), H3 (middle inset), and Flu B (bottom inset). See ED Fig. 4 for an equivalent graph for CR-6261 binding to H1 and H9.

Beyond single mutational effects, it remains unknown whether there are correlated effects or strong trade-offs between binding to different antigens (pleiotropy), or non-additive interactions between mutations (epistasis). Such epistatic and pleiotropic effects can constrain the mutational pathways accessible under selection, as has been observed for other proteins^14–22^. Epistasis in antibody-antigen interactions remains significantly understudied^23–25^, and most deep mutational scanning studies have focused on antigens^26–28^. In contrast to typical protein evolution, antibody affinity maturation proceeds by discrete rounds of mutation and selection^12^, typically with more than one nucleotide mutation occurring between selective rounds^29^. In addition, antibodies are inherently mutationally tolerant^25,30,31^, and bnAbs tend to have many more mutations than specific antibodies^2,32^, generating opportunities for interactions that scale combinatorially. Thus, if epistatic and pleiotropic constraints exist for antibodies, they could affect the likelihood of producing bnAbs under different antigen selection regimes^24^ and may account for the low frequencies of bnAbs in natural repertoires^1^. Characterizing the prevalence of these constraints on bnAb evolution will provide valuable insight for designing optimal vaccination strategies^33,34^.

To date, studies of antibody binding have been limited to small numbers of individual sequences, deep mutational scans of single mutations, and mutagenesis of small regions ^24,25,31,35–40^, due in part to practical constraints on library scale and the throughput of affinity assays. This has limited our ability to comprehensively characterize binding landscapes for naturally isolated bnAbs, which often involve many mutations spanning framework (FW) and complementarity-determining regions (CDR)^1,2,32^. Here, we overcome these challenges by generating combinatorially complete libraries of up to ~10^5^ antibody sequences and assaying their binding affinities in a high-throughput yeast-display system^35^. Specifically, we made all combinations of 16 mutations from CR-9114 (65,536 variants) and all combinations of 11 mutations from CR-6261 (2,048 variants). These libraries include all heavy-chain mutations in these antibodies, except select mutations distant from the paratope (Fig. 1a, and see SI). Both antibodies engage antigens solely through their heavy-chain regions^4,5^, and thus are well-suited for yeast display as single-chain variable fragments (see SI)^41^.

We use the Tite-Seq method^35^, which integrates flow cytometry and sequencing (ED Fig. 2b), to assay equilibrium binding affinities of each sequence in these libraries against select antigens that span the breadth of binding for each antibody (Fig 1b). For CR-6261, we chose two divergent group 1 HA subtypes (H1 and H9), while for CR-9114, we chose the three highly divergent subtypes present in the vaccine (H1 from group 1, H3 from group 2, and influenza B)^3^. Inferred affinities outside our titration boundaries (10^−11^ – 10^−6^ M for H3 and influenza B, 10^−12^ – 10^−7^ M for H1 and H9) are pinned to the boundary, as deviations beyond these boundaries are likely not physiologically relevant^42^. Antibody expression is not strongly impacted by sequence identity, although some mutations have modest effects that may be inversely correlated with their effect on affinity (ED Fig. 3). Affinities obtained by Tite-Seq are reproducible across biological triplicates (ED Fig. 2c,e; average standard error of 0.047 −logK_D_ units across antibody-antigen pairs) and are highly accurate as verified for select variants by isogenic flow cytometry (ED Fig. 2d,f) and by solution-based affinity measurements made by others^3,4,13,24^.

We begin by examining the distribution of binding affinities across antigens for each antibody library (Fig. 1). We observe that most CR-9114 variants have measurable affinity to H1 (97%), fewer to H3 (11%), and still fewer to influenza B (0.3%) (Fig. 1c,d). For H1, only a few mutations are needed to improve from the germline affinity. In contrast, variants are not able to bind H3 unless they have several more mutations, and many more for influenza B. This hierarchical structure is in striking contrast to the CR-6261 library, in which most variants can bind both antigens (92% for H1, 81% for H9), variants have a similar K_D_ distribution, and many variants display intermediate affinity to both antigens (Fig. 1e,f). To visualize how genotypes give rise to the hierarchical structure of CR-9114 binding affinities, we represent the binding affinities for H1 as a force-directed graph. Here, each variant is a node connected to its 16 single-mutation neighbors, with edge weights inversely proportional to the change in H1 binding affinity, such that variants with similar genotype and K_D_ tend to form clusters (Fig. 1g, ED Fig. 4). Coloring this genotype-to-phenotype map by the −logK_D_ to each of the three antigens, we see that sequences that bind H3 and influenza B are highly localized and overlapping, meaning that they share specific mutations. Thus, while many CR-9114 variants strongly bind H1, only a specific subset bind multiple antigens.

To dissect how mutations drive the structure of these binding landscapes, we next infer specific mutational effects. We first log-transform binding affinities to produce free energy changes, which should combine additively under the natural null expectation^43,44^. We then define a linear model with single mutational effects and interaction terms up to a specified order (defined relative to the unmutated germline sequence, see SI for alternatives), and fit coefficients by ordinary least squares regression (see SI for models with nonlinear transformations). We use cross-validation to identify the maximal order of interaction for each antigen and report coefficients at each order from these best-fitting models (CR-9114: 5^th^ order for H1, 4^th^ for H3, 1^st^ for influenza B; CR-6261: 4^th^ order for H1 and H9; see SI). We note that the maximum order of interactions is affected by our inference power, particularly by the number of sequences with appreciable binding, and so we interpret these models as showing strong evidence of epistasis at least up to the order indicated. We also explored inferring epistasis up to full order using Walsh-Hadamard transformations; results are qualitatively similar but less conservative than cross-validated regression (see SI).

Examining the effect of individual mutations on the germline background (Fig. 2a,b), we observe several mutations that enhance binding to all antigens (e.g. S83F for CR-9114), and mutations that confer trade-offs for binding distinct antigens (e.g. F30S in CR-9114 reduces affinity for H1 but enhances affinity for influenza B). Generally, large-effect mutations are at sites that contact HA^4,5^ (Fig. 2c, ED Fig. 5). Consistent with prior biochemical and structural work, mutations essential for CR-9114 breadth are spread throughout FW3 and the CDRs, forming hydrophobic contacts and hydrogen bonds with residues in the conserved HA stem epitope^4^. We observe three specific mutations that are required for binding to H3 (present at over 90% frequency in the set of binding sequences), and eight specific mutations that are required for binding to influenza B. Many of these breadth-conferring mutations are absent in CR-6261, particularly those in CDR2^4,5^. Notably, these sets of required mutations in CR-9114 exhibit a nested structure: mutations beneficial for H1 are required for H3, and mutations required for H3 are required for influenza B, giving rise to the hierarchical structure of the binding landscape (Fig. 1c).

**Figure 2:**
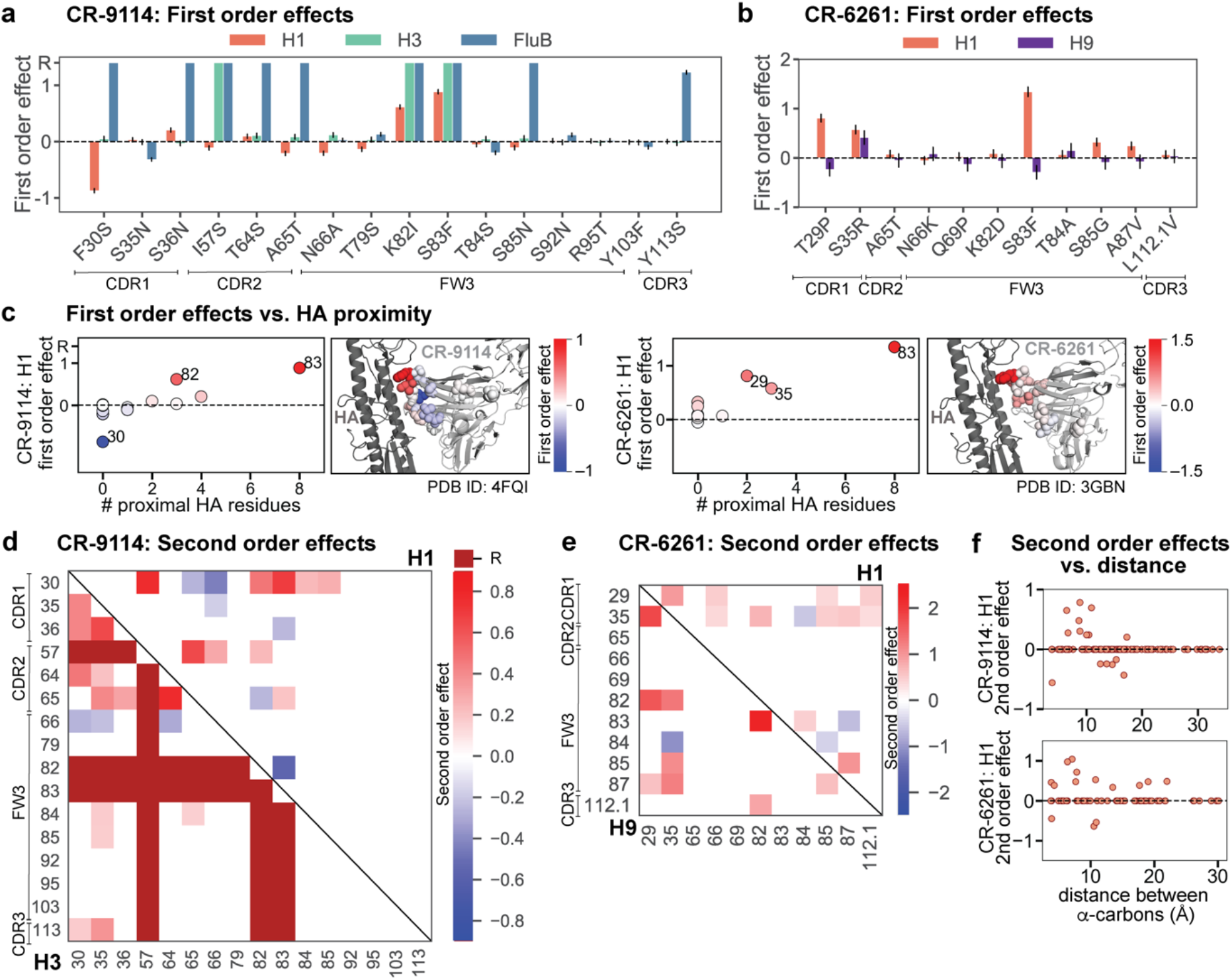
Linear effects and pairwise epistasis. **a, b** First order effects inferred in best-fitting epistatic interaction models for **(a)** CR-9114 and **(b)** CR-6261. Mutations required for binding (present at over 90% frequency in binding sequences) have effect sizes denoted as “R” and are removed from inference. Error bars indicate standard error. **c,** First order effects for each site plotted against the number of HA residues within 6 Å (left, CR-9114; right, CR-6261). Top three sites are labeled. Cocrystal structures are also shown; mutations are colored by first-order effect size. See ED Fig. 5a for equivalent plot for CR-9114 with H3 and CR-6261 with H9. **d,** Significant second-order epistatic interaction coefficients for CR-9114 mutations (bottom left, H3; top right, H1). Interactions involving required mutations are shown in dark red. **e,** Significant second order coefficients for CR-6261 mutations (bottom left, H9; top right, H1). **f,** Second-order coefficients for H1 −logK_D_ plotted against the distance between the respective α-carbons in the crystal structures. See ED Fig. 5b for equivalent plot for CR-9114 with H3 and CR-6261 with H9. Significance in **d**, **e** indicates Bonferroni-corrected *p*-value < 0.05, see SI.

Beyond these exceptionally synergistic interactions between required mutations, we find that epistasis is widespread, accounting for 18–33 percent of explained variance depending on the antibody-antigen pair (except influenza B, see SI). Pairwise interactions are dominated by a few mutations (e.g. F30S for CR-9114 and S35R for CR-6261) that exhibit many interactions, both positive and negative, with other mutations (Fig. 2d,e). Overall, mutations with strong pairwise interactions tend to be close in the crystal structure (Fig. 2f, ED Fig. 5)^4,5^.

Our dataset also allows us to resolve higher-order epistasis. In addition to the required mutations, our models identify numerous strong third to fifth order interactions, with a subset of mutations participating in many mutual interactions at all orders. For CR-9114 binding to H1, this subset consists of five mutations, distributed across three different regions of the heavy chain (Fig. 3a,b). Some of these mutations likely generate (K82I, S83F) or abrogate (F30S) contacts to HA, and others (I57S, A65T) may indirectly impact HA binding by reorienting contact residues in CDR2^4^. Within this set of five residues, we first illustrate two examples of third-order epistasis by grouping all sequences by their genotypes only at these five sites (Fig. 3c). Intriguingly, some mutations that are deleterious in the germline background (‘–’ annotations) are beneficial in doubly-mutated backgrounds (‘+’ annotations). For example, mutation F30S is significantly less deleterious in backgrounds with S83F than in the germline background, suggesting that new hydrophobic contacts in FW3 may be able to compensate for the potential loss of contacts in CDR1. Yet F30S unexpectedly becomes beneficial after an additional mutation I57S in CDR2, indicating more complex interactions between flexible CDR and FW loop regions (Fig. 3b,c)^4^.

**Figure 3:**
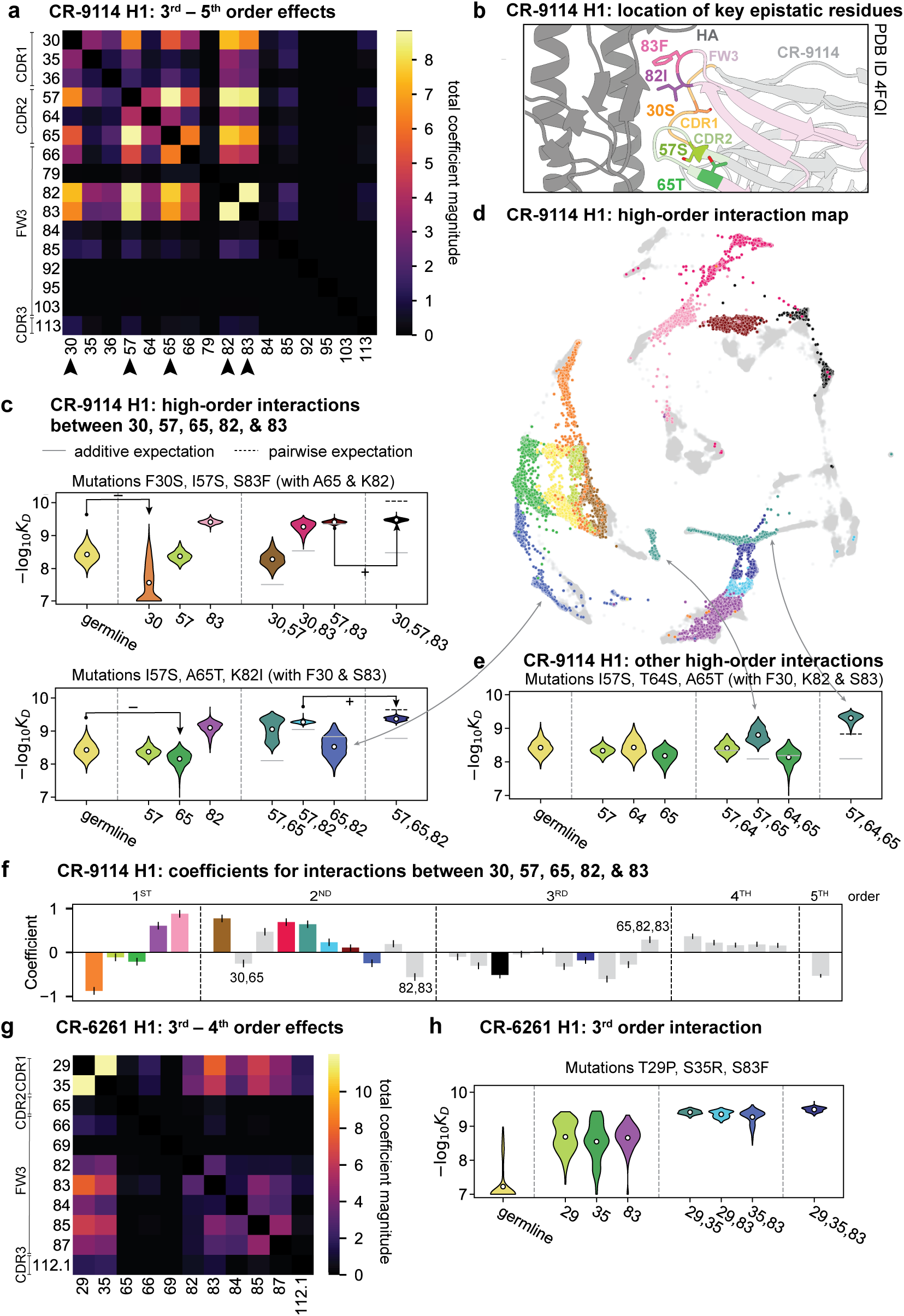
High-order epistasis. **a**, Total higher-order epistatic contributions of CR-9114 mutation pairs for binding H1. Color bar indicates the sum of absolute values of significant higher-order interaction coefficients involving each pair of mutations; key epistatic residues indicated by arrows. See ED Fig. 6f for equivalent figure with H3. **b**, Location of key epistatic residues in the CR-9114–HA co-crystal structure colored by region. **c**, – logK_D_ distributions for genotypes grouped by their identity at the five residues indicated in **a, b,** with means indicated as white dots (N=8,192 genotypes per violin). Annotations indicate notable deleterious (‘−’) and beneficial (‘+’) mutational effects. **d,** CR-9114 force-directed graph from Fig. 1g, colored as in **c** by the genotype at the five sites indicated in **a, b**. Genotypes not shown in **c** are shown in light grey. See ED Fig. 6a for graph colored by all 32 5-site genotypes. **e**, Third-order interaction involving site 64 accounts for distinct clusters (teal) corresponding to genotypes with mutations 57 and 65 in **d**. Colors correspond to mutation groups in **c, d** (N=4,096 genotypes per violin). **Figure 3, continued**: **f**, Epistatic interaction coefficients among the five key sites from **a, b**. Colors for certain groups as in **c, d**; other groups denoted in gray, with notable terms labeled. **g,** Total significant epistatic contributions of CR-6261 mutation pairs for binding H1, as in **a**. **h**, Third-order interaction for CR-6261 H1 binding between mutations T29P, S35R, and S83F (N=256 genotypes per violin). Significance in **a**, **g** indicates Bonferroni-corrected *p*-value < 0.05, see SI.

To see how these high-order interactions drive the overall structure of the binding affinity landscape, we return to the force-directed graph, now colored by genotype at these five key sites (Fig. 3d; only points corresponding to genotypes shown in Fig. 3c are colored). We see that these five sites largely determine the overall structure of the map: points of the same color tend to cluster together, despite varying in their genotypes at the other 11 sites. However, we observe that interactions with other mutations do exist, as evidenced by separate clusters with the same color (e.g. the two clusters in teal for 57,65 are distinguished by a positive third-order interaction with site 64, Fig. 3e). These patterns are not confined to the genotypes shown in Fig. 3c; if we color all 32 possible genotypes at the five key sites, we observe the same general patterns (ED Fig. 6; an interactive data browser for exploring these patterns of epistasis is available <u>here</u>). Interactions between these five sites are also enriched for significant epistatic coefficients (p < 10^−3^; 26 of 31 possible terms are significant, compared to an average of 4 terms among all sets of five sites, ED Fig. 6), including the 5^th^ order interaction between all five residues (Fig. 3f). Remarkably, these five mutations underlie significant high-order epistasis for other antigens as well: all five are either required for binding or participate extensively in interactions for H3 and influenza B (ED Fig. 6f).

Higher-order epistasis in CR-6261 is similarly dominated by a subset of mutations in CDR1 and FW3, at identical or neighboring positions as some key sites for CR-9114 (Fig. 3g). These mutations exhibit strong diminishing returns epistasis at third and fourth order, counteracting their synergistic pairwise effects, in a similar manner across both antigens (Fig. 3h, ED Fig. 7). Many fourth-order combinations of these mutations display interaction coefficients of similar magnitude (ED Fig. 7b), though they may be signatures of even higher-order interactions that we are underpowered to infer.

A common approach to quantify how epistasis constrains mutational trajectories is to count the number of “uphill” paths (i.e. where affinity improves at every mutational step from the germline to the somatic sequence). We find that only a small fraction of potential paths are uphill (0.00005% +/− 0.00004% for CR-9114 binding H1, and 0.2% +/− 0.04% for CR-6261 binding H1, as estimated by bootstrap, see SI). However, we note that for all antibody-antigen combinations, the somatic sequence is not the global maximum of the landscape (the best-binding sequence) and some mutations have deleterious effects on average. Hence, strictly uphill paths are only possible due to sign epistasis, where normally deleterious mutations have beneficial effects in specific genetic backgrounds.

Overall, we see that mutational effects and interactions between them explain the affinity landscapes we observe. For CR-9114, binding affinity to H1 can be achieved through different sets of few mutations with complex interactions. In contrast, a specific set of many mutations with strong synergistic interactions is required to bind H3, and to an even greater extent, influenza B, giving rise to the landscape's hierarchical structure (Fig. 1c). For CR-6261, the higher-order interactions are more similar between H1 and H9, which is consistent with the more correlated patterns of binding affinities between these two antigens (Fig. 1e).

The hierarchical nature of the CR-9114 landscape suggests that this lineage developed affinity to each antigen sequentially. Considering the maximum −logK_D_ achieved by sequences with a given number of mutations (a proxy for time), we see that improvements in H1 binding can be realized early on, whereas improvements in H3 binding are not possible until later, and even later for influenza B (Fig. 4a). In fact, the nested structure of affinity-enhancing mutations forces improvements in binding affinity to occur sequentially. If selection pressures were also experienced in this sequence, mutations that improve binding to the current antigen would lead to the genotypes required to begin improving binding to the next. Indeed, we find that for CR-9114, there are more uphill paths leading to the somatic sequence if selection acts first on binding to H1 and later to H3 and influenza B (Fig. 4c). In contrast, for CR-6261, improvements in binding can occur early on for both antigens (Fig. 4b) and the number of uphill paths is more similar across single-antigen and sequential selection pressures (Fig. 4d).

**Figure 4:**
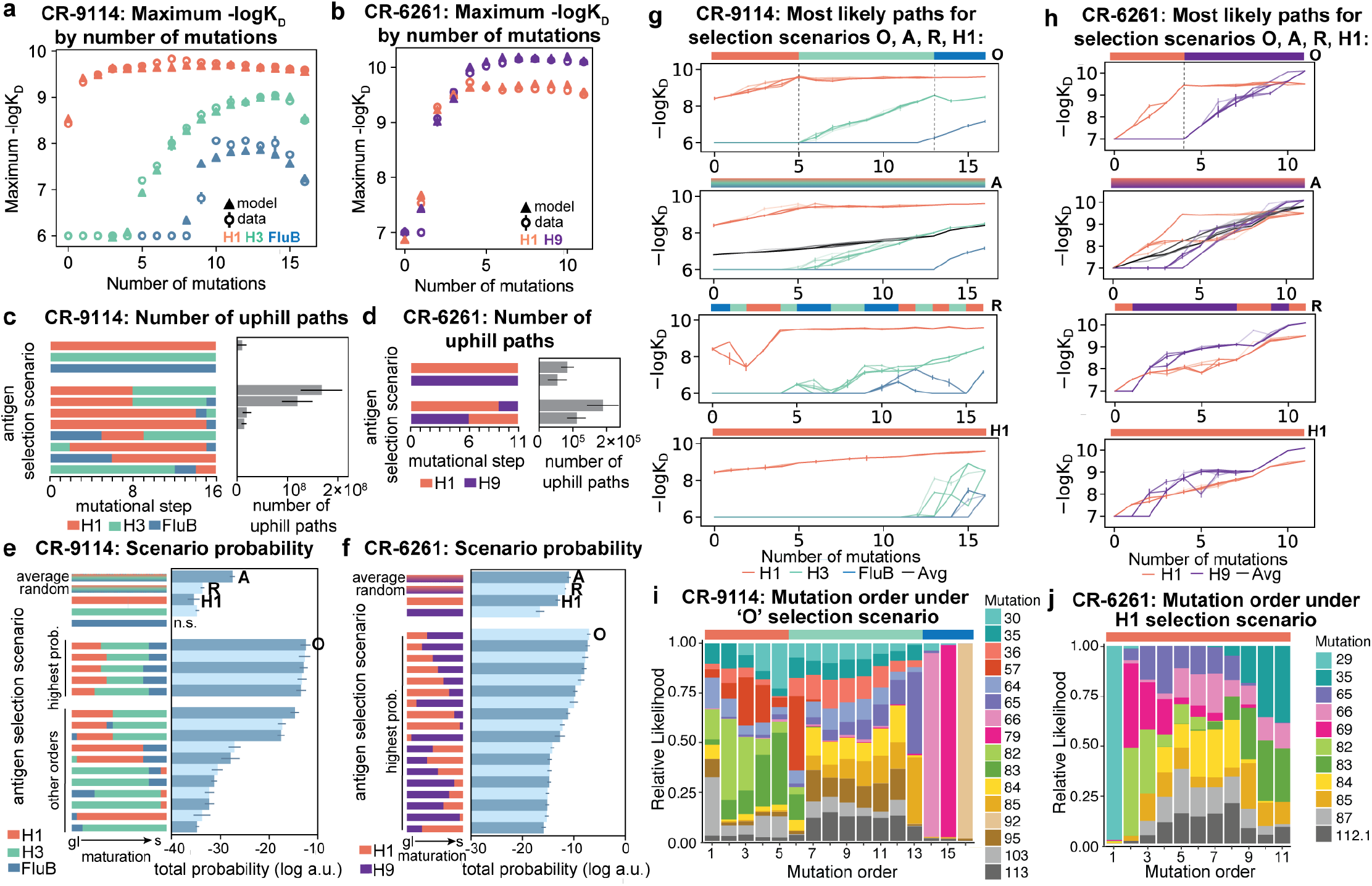
Antigen selection scenarios and likely mutational pathways. **a, b,** Maximum binding affinity achievable for sequences with a given number of mutations. For each antigen for (**a**) CR-9114 and (**b**) CR-6261, the maximum observed (circles) and model-predicted (triangles) affinity for each number of somatic mutations is shown. **c, d,** Total number of ‘uphill’ paths for select antigen selection scenarios (colored bars) for (**c**) CR-9114 and (**d**) CR-6261. Error bars indicate standard error obtained through bootstrap. **e, f,** Total log probability (in arbitrary units) of mutational trajectories from germline to somatic sequence under different antigen selection scenarios, in a moderate selection model (see ED Fig 8a,b for other models). Error bars indicate standard error obtained through bootstrap. **g, h**, 25 most likely paths for (**g**) CR-9114 and (**h**) CR-6261, from select scenarios in **e, f**; −logK_D_ plotted for each antigen. For the random mixed scenario (‘R’), a representative case is shown. **i, j,** Probability of mutation order under antigen selection scenario ‘O’ for CR-9114 (**i**) and ‘H1’ for CR-6261 (**j**). Selection scenarios are as in **e, f** and shown in colored bar at top; the total probability (through all possible paths) for each mutation to occur at each mutational step is shown as stacked colored bars. See ED Fig. 8 for additional selection scenarios.

To compare antigen selection scenarios more generally, we developed a framework that evaluates the total probability of all possible mutational pathways from germline to somatic, under an array of antigen selection scenarios (individual, sequential, and mixed). Our framework assumes that the probability of any mutational step is higher if −logK_D_ increases, but does not necessarily forbid neutral or deleterious steps; we evaluate a variety of specific forms of this step probability and find that our major results are consistent (ED Fig. 8a, see SI). Mixed antigen regimes approximate exposure to a cocktail of antigens. We model these with two approaches: (1) “average”, using the average −logK_D_ across all antigens, and (2) “random,” using −logK_D_ for a randomly selected antigen at each step (note that using the maximum −logK_D_ across antigens would always be trivially favored)^6^. While these models simplify the complexities of affinity maturation *in vivo*^12^, especially in how affinity relates to B cell lineage dynamics, they provide insight into the relative probabilities of mutational paths under distinct antigen selection scenarios.

Again we find that the vast majority of likely antigen selection scenarios for CR-9114 involve first H1, followed by H3, followed by influenza B (Fig. 4e, ED Fig. 8b). These results are underscored by examining improvement in −logK_D_ along the most likely mutational paths for each scenario (Fig. 4g): in the optimal sequential scenario, −logK_D_ can improve substantially for each antigen in turn, while in an H1-only scenario, the improvements in H1 binding at each step are much more gradual, reducing the likelihood. The average mixed scenario shows qualitatively similar paths to the optimal sequential scenario, although with lower overall probability. In the random mixed scenario, even the best pathways are often unable to improve affinity to the randomly selected antigen, and affinity to antigens not under selection often declines, making these scenarios much less likely.

Given the optimal sequential selection scenario, the vast majority of genotypes are unlikely evolutionary intermediates to the somatic sequence (ED Fig. 8d). We visualize the impact of epistasis on mutational order by considering the probability of each mutation to occur at each mutational step (Fig. 4i; ED Fig. 8e,f). The three antigen exposure epochs exhibit clear differences in favored mutations. Mutations I57S, K82I, and S83F must occur early, due to their strong synergistic interactions for all three antigens. In addition, we see that F30S is unlikely to happen very early (due to its sign epistasis under H1 selection) as well as unlikely to happen very late (due to its strong benefit under influenza B selection).

In contrast, for CR-6261, all selection scenarios have relatively similar likelihood (Fig. 4f, ED Fig. 8c). Among sequential scenarios, however, those beginning with H1 are more likely than those beginning with H9, as the first two mutational steps can improve affinity to H1 more than H9, and mutations late in maturation can improve affinity to H9 more than H1 (Figs. 1f, 4b). Still, unlike CR-9114, in both single antigen and mixed scenarios, there are many likely paths that continually improve in binding to both antigens (Fig. 4h). Initially the order of mutations is highly constrained due to strong synergistic epistasis, and differences between selection scenarios reflect differences in mutational effects between antigens (Fig. 4h, ED Fig. 8g,h).

Overall, we find that evolutionary pathways to bnAbs can be highly contingent on epistatic and pleiotropic effects of mutations. Specifically, the acquisition of breadth for CR-9114 is extremely constrained and is likely to have occurred through exposure to diverse antigens in a specific order, due to the structure of correlations and interactions between mutational effects. In contrast, CR-6261 could have acquired affinity to H1 and H9 in a continuous and simultaneous manner, perhaps because these antigens are more similar; since H9 is not a commonly circulating strain, this breadth may well have been acquired by chance^24^. Though we cannot conclusively determine which antigens were involved in the selection of these antibodies *in vivo*, the diverse HA subtypes discussed here capture variation representative of circulating influenza strains and thus serve as useful probes of varying levels of breadth^1^. Further, we note that the likelihood of pathways is conditioned on ending at the exact somatic sequence. Indeed, we observe that not all of the observed mutations are required to confer broad affinity, and future work is needed to explore what alternative pathways to breadth might be accessible through other mutations.

The landscapes characterized here are among the largest combinatorially complete collections of mutations published to date. In some respects, our observations of high-order interactions are consistent with earlier work in other proteins. In particular, epistasis has been found to affect function and constrain evolutionarily accessible pathways across functionally and structurally distinct proteins^14–22^. Further, pairwise and high-order epistasis appear to be common features of binding interfaces, such as enzyme-substrate and receptor-ligand interactions^14–17,19,20^, and interacting mutations are often spaced in both sequence and structure, underscoring the complexity of protein-protein interfaces^17,23,25,45,46^. On the other hand, the strongly synergistic, nested mutations crucial for CR-9114 breadth are unusual, perhaps due to the nature of antibody-antigen interfaces or to the unique dynamics of affinity maturation^12^. Together, these observations suggest that interactions between multiple mutations, such as those we characterize here, could play a substantial role in affinity maturation and may contribute to the rarity of bnAbs in natural repertoires.

Our findings provide molecular insight into the emerging picture of how selection can elicit broad affinity, illustrated by a substantial recent body of work ranging from *in vivo* experimental approaches^7,8^ to quantitative modeling of immune system dynamics^6,47–50^. These diverse studies often find that mixed-antigen regimens are less effective than sequential regimens at eliciting bnAbs. Our results demonstrate that, at least in part, this may be due to the intrinsic structure of the mutational landscape, defined by the complex interactions of mutational effects across antigens. With more studies of binding landscapes for diverse antibodies, we could better understand how such features generalize between different germline sequences, somatic mutation profiles, and antigen molecules. These insights will be essential for leveraging germline sequence data and antigen exposure information to predict, design, and elicit bnAbs for therapeutic and immunization applications.

## ACKNOWLEDGEMENTS

We thank Rhys Adams for helpful discussion of the Tite-Seq experiments, Zach Niziolek for assistance with flow cytometry, Kevin McCarthy for help with antigen production, Matt Melissa for help acquiring strains and protocols, and Tyler Starr and members of the Denic, Gaudet, and Wittrup labs for help with experimental protocols. We also thank Jesse Bloom, Andrew Murray, and Michael Laub for helpful discussion and members of the Desai lab for comments on the manuscript. A.M.P. acknowledges support from the Howard Hughes Medical Institute Hanna H. Gray Postdoctoral Fellowship Program. K.R.L. acknowledges support from the Fannie & John Hertz Foundation Graduate Fellowship Award and the NSF Graduate Research Fellowship Program. T.D. acknowledges support from the Human Sciences Frontier Program. J.C. acknowledges support from the NSF Graduate Research Fellowship Program. M.S.J. acknowledges support from the NSF Graduate Research Fellowship Program. Work in Paris was funded by the European Research Council COG 724208. M.M.D. acknowledges support from grant PHY-1914916 from the NSF and grant GM104239 from the NIH. The computations in this paper were run on the FASRC Cannon cluster supported by the FAS Division of Science Research Computing Group at Harvard University.

## AUTHOR CONTRIBUTIONS

A.M.P., K.R.L, I.C., M.M.D, A.W., and T.M. conceived the project; A.M.P., K.R.L., and A.M. generated the yeast display libraries; A.M.P., K.R.L., A.M., J.C., and I.C. conducted experiments and binding affinity measurements; A.M.P., K.R.L., A.M., and T.D. developed inference methods and conducted statistical analysis; M.S.J. developed the interactive data browser; A.M.P., K.R.L., M.M.D., A.W., and T.M. wrote the paper.

Raw and processed data are available in an interactive data browser here: https://yodabrowser.netlify.app/yoda_browser/

---

**ED Figure 1:**
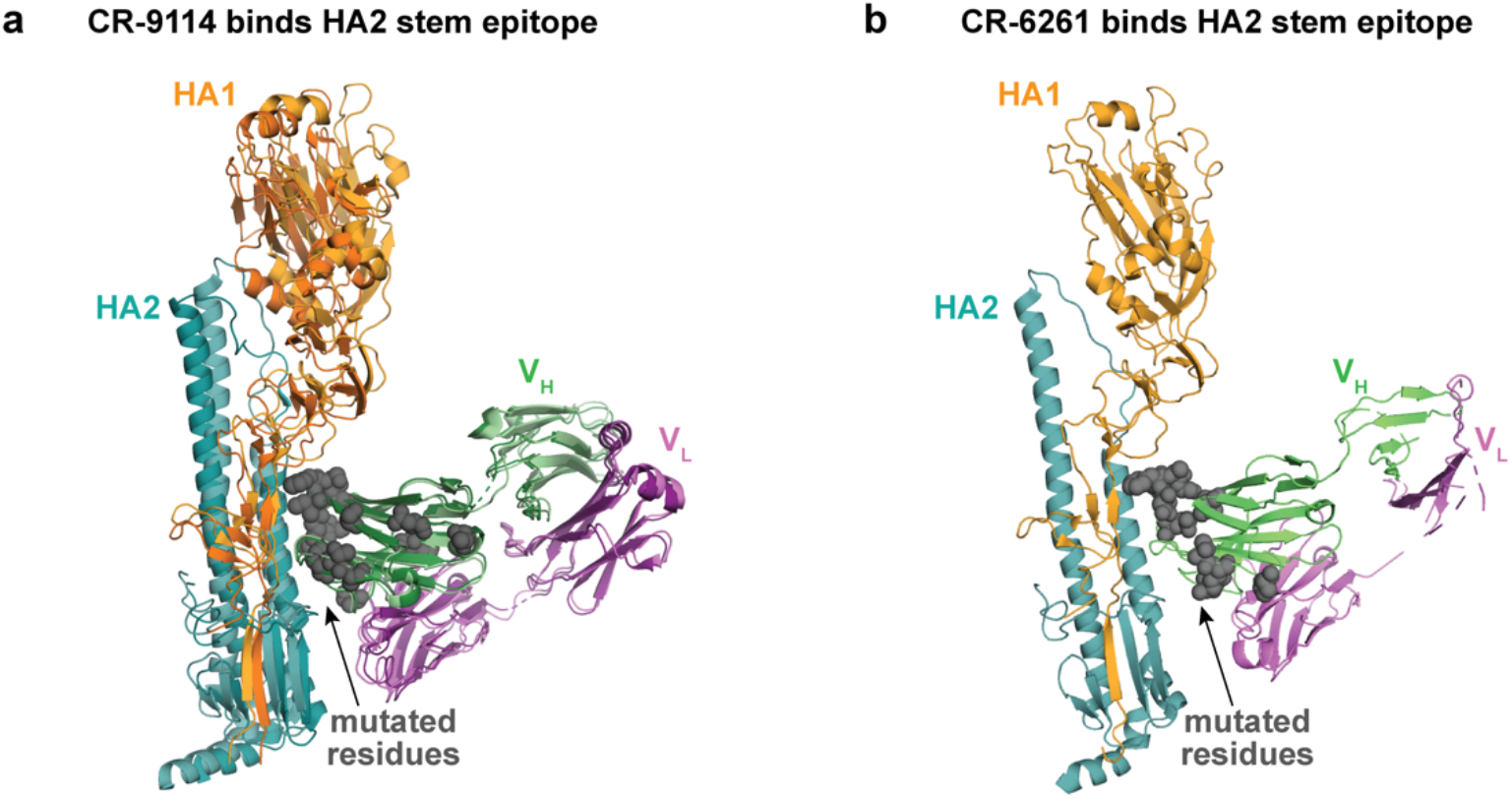
Co-crystal structures. **a**, Alignment of co-crystal structure of CR-9114 with H5 (light hues; PDB ID 4FQI^4^) and CR-9114 with H3 (dark hues; PDB ID 4FQY^4^). Mutated residues shown as gray spheres. **b**, Co-crystal structure of CR-6261 with H1 (PDB ID 3GBN^5^); mutated residues shown as gray spheres.

**ED Figure 2:**
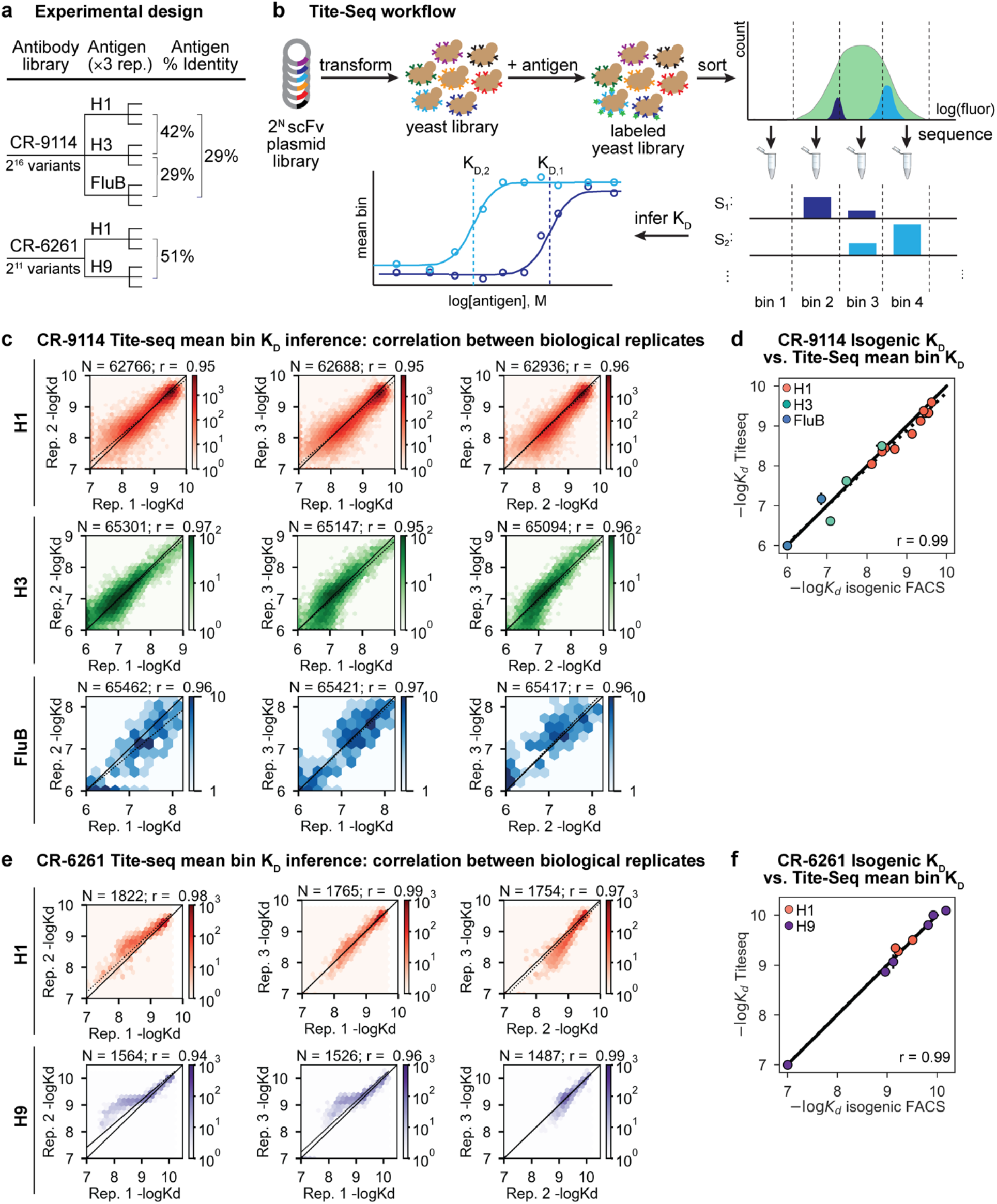
Data quality. **a,** Experimental design. **b,** Tite-Seq assay. Surface display single-chain variable fragment (scFv) libraries are transformed into yeast and labeled with fluorescent antigen, followed by FACS into bins and sequencing. Dissociation constants are inferred from changes in mean bin fluorescence across 12 antigen concentrations, see SI. **c, e**, Correlation of (**c**) CR-9114 and (**e**) CR-6261 K_D_ measurements between biological replicates. **d, f,** Validation of (**d**) CR-9114 and (**f**) CR-6261 Tite-Seq K_D_ measurements by isogenic flow cytometry measurements for a subset of variants and antigens.

**ED Figure 3:**
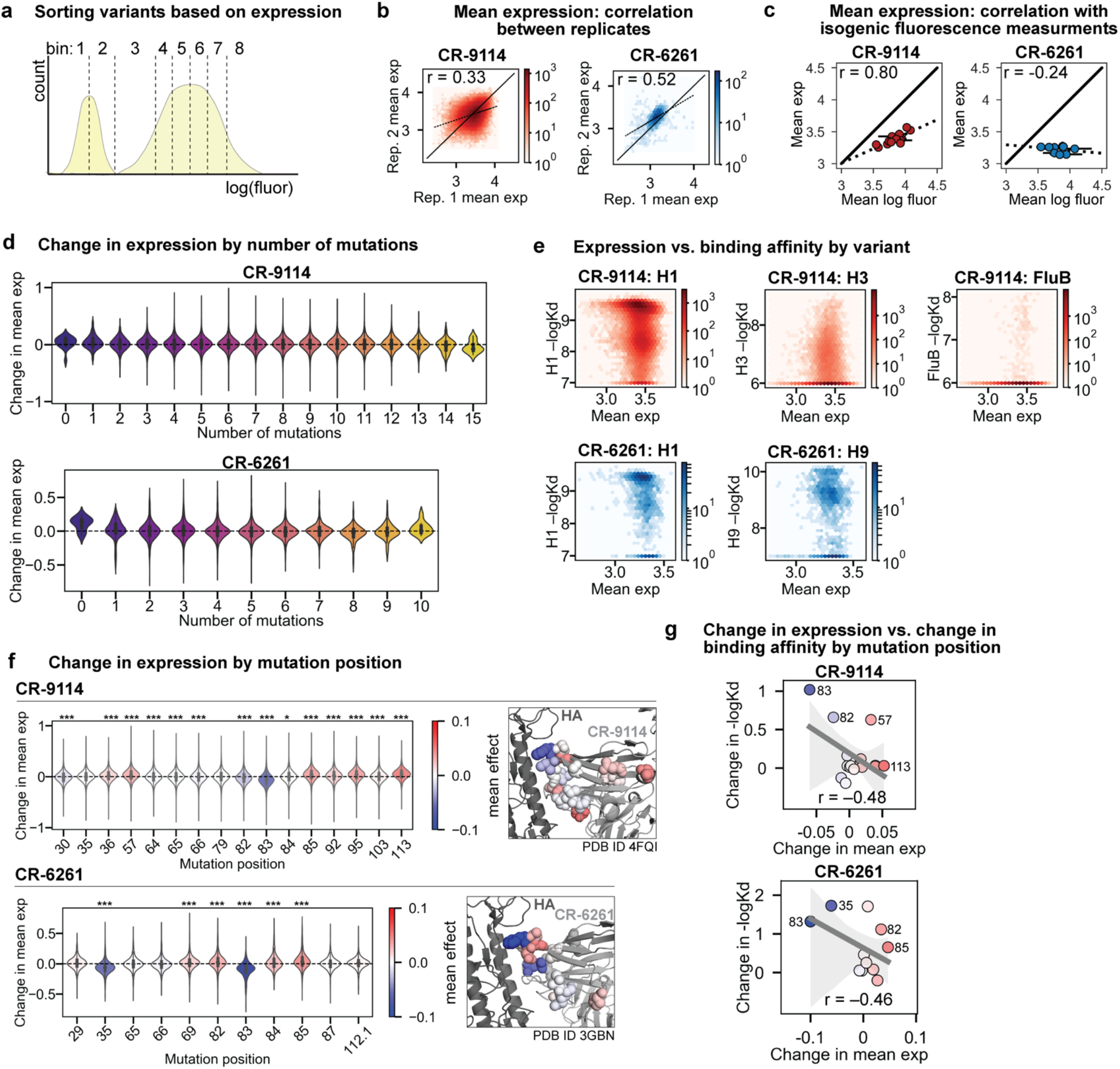
Expression of antibody libraries. **a**, Expression gates were drawn such that each of 8 bins included 12.5% of the antibody library. Expression is calculated as mean log fluorescence of each sequence across bins (see SI). **b**, Correlation of expression across biological replicates for CR-9114 library (left, red) and CR-6261 library (right, blue). **c**, Correlation between Tite-Seq mean expression and isogenic expression fluorescence for select CR-9114 (left, red) and CR-6261 (right, blue) variants. **d**, Change in expression upon mutation for a given number of background somatic mutations. **e,** Correlation between mean expression and −logK_D_. Average values across biological replicates (N_−logKD_ = 3; N_exp_ ≥ 6) are plotted. **f**, Change in expression upon mutation at a specific site. Violin plots (left) and residues in co-crystal structure (right) are colored by mean change in expression for each site. Asterisks above violins indicate *p*-values for two-sided *t*-test between the distribution means and zero (p < 0.01 (*), < 0.001 (**), < 0.0001 (***)). **g**, Correlation between mean change in expression and mean change in −logK_D_ (summed across all antigens) by mutation position. Select mutations with large impacts on expression and −logK_D_ are labeled; all points are colored by mean change in expression, as in **f**. Dark gray line indicates best-fit linear regression (95% confidence intervals in light gray).

**ED Figure 4:**
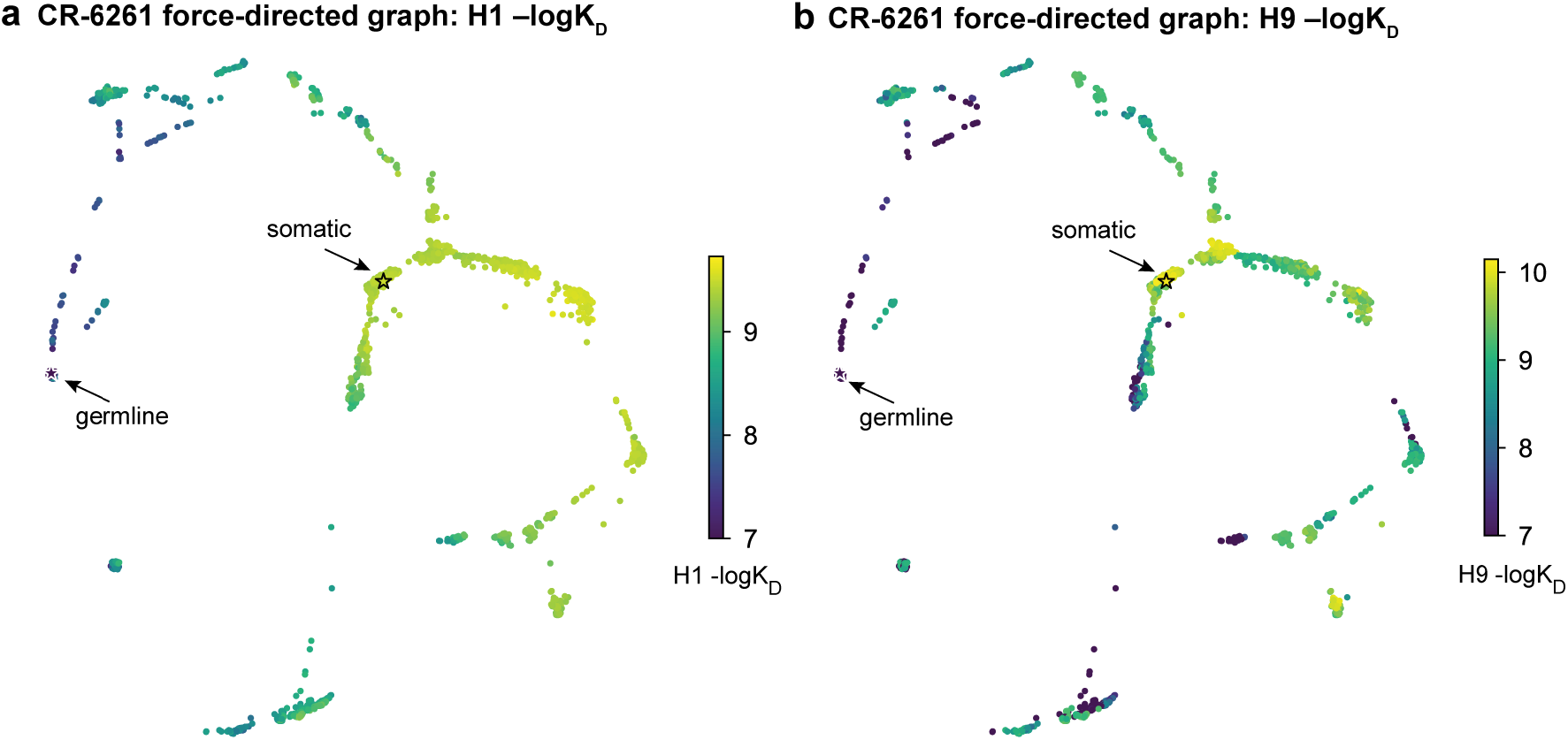
Force-directed graphs. **a, b,** Force-directed graph for CR-6261 H1 −logK_D_, as in Fig. 1g. Nodes are colored by binding affinity to (**a**) H1 and (**b**) H9.

**ED Figure 5:**
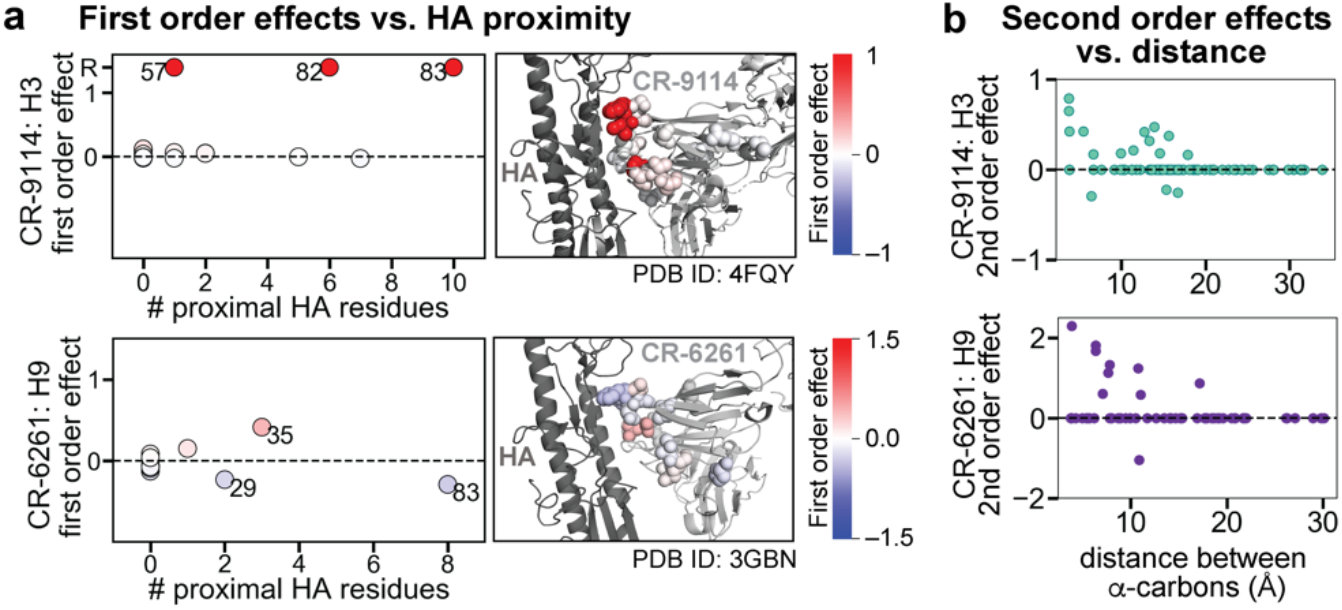
Structural context of first and second order effects. **a**, Left: first order effects for each site, colored by effect size and plotted against the number of antigen residues within 6 Å (top, CR-9114 with H3; bottom, CR-6261 with H9); Right: cocrystal structures with mutation sites colored by first order effects, as in Fig. 2c. **b,** Second-order coefficients for CR-9114 (top) and CR-6261 (bottom) plotted against the distance between the respective α-carbons in the crystal structures, as in Fig. 2f.

**ED Figure 6:**
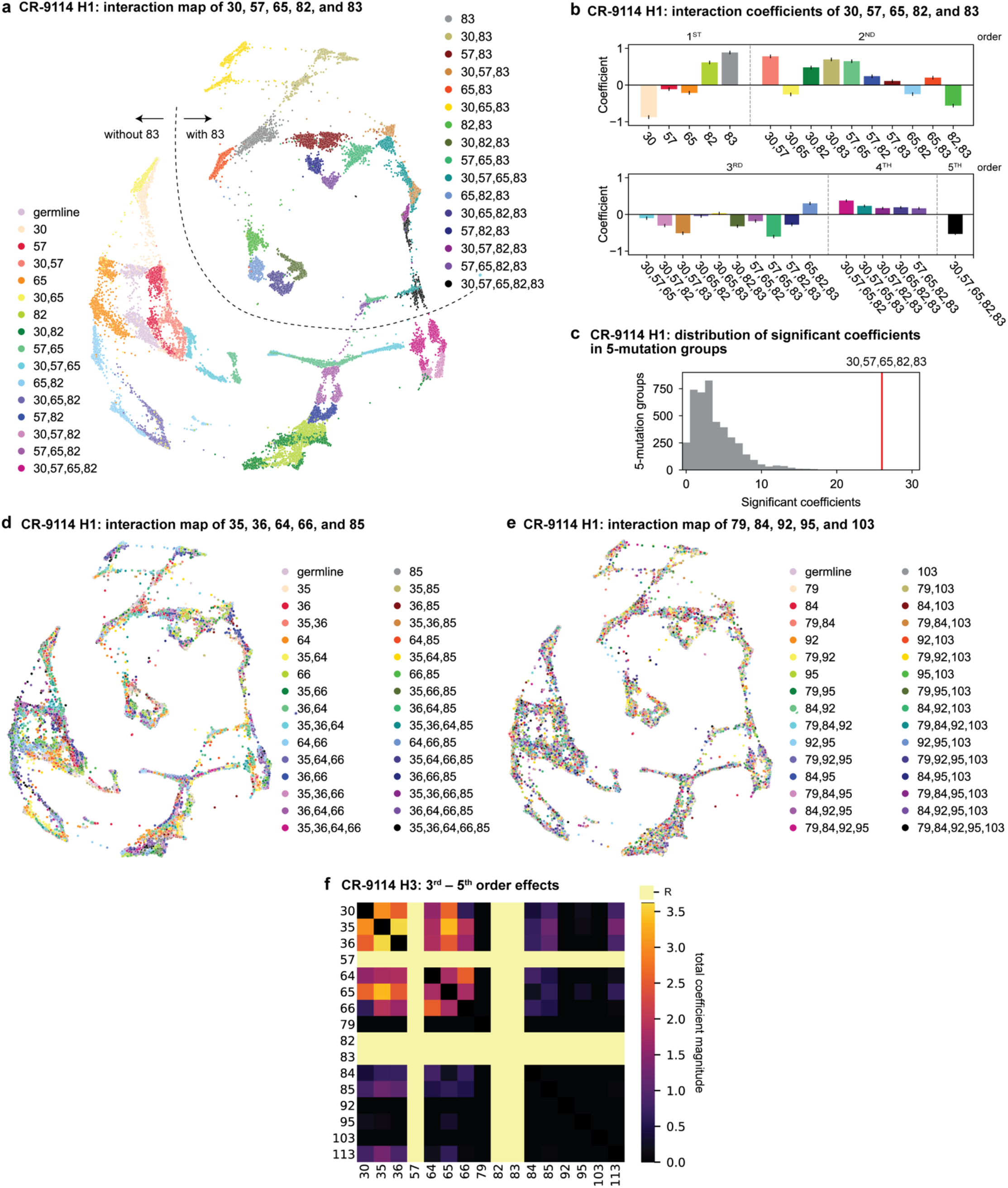
High-order interactions for CR-9114. **a**, CR-9114 force-directed graph, as in Fig. 3d, colored by mutation groups of sites 30, 57, 65, 82, and 83 (32 total groups). The dashed line emphasizes the observed separation of genotypes with S83 (upper left) from those with S83F (lower right). **b**, Coefficients for terms in the epistatic interaction model corresponding to mutation groups of sites 30, 57, 65, 82, and 83 (31 total groups, excluding the germline), colored according to **a** and grouped by order. Error bars indicate standard error. **ED Figure 6, continued: c**, Distribution of the number of significant coefficients for mutation groups in every possible set of 5 sites chosen from the 16 sites (up to 31 terms for each group, for 4,368 groups). The group illustrated in **a, b** is shown in red (26 significant terms, empirical p-value < 10^−3^). **d**, CR-9114 force-directed graph, colored by mutation groups of a different set of 5 sites with fewer strong epistatic interactions (35, 36, 64, 66, and 85). **e**, CR-9114 force-directed graph, colored by mutation groups of a different set of 5 sites with no strong linear contributions or epistatic interactions (79, 84, 92, 95, and 103). **f,** Higher-order significant epistatic contributions of CR-9114 mutation pairs, as in Fig. 3b, for binding H3. Light yellow columns indicate required mutations (sites 57, 82, and 83). Significance in **c**, **f** indicates Bonferroni-corrected *p*-value < 0.05, see SI.

**ED Figure 7:**
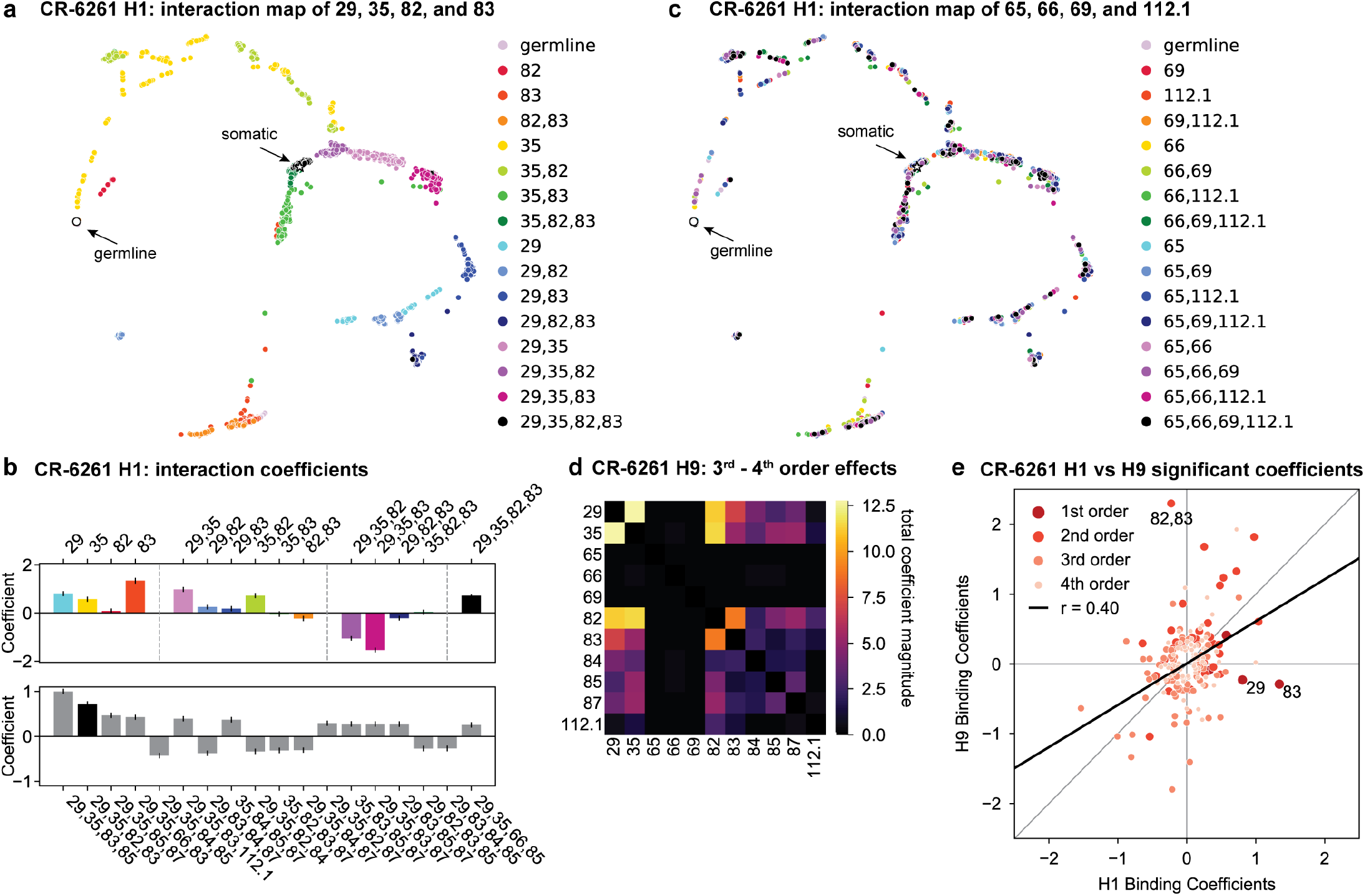
High-order interactions for CR-6261. **a**, CR-6261 force-directed graph, as in ED Fig. 4, colored by mutation groups of sites 29, 35, 82, and 83 (16 total groups). **b**, Top, coefficients for terms in the epistatic interaction model corresponding to the mutation groups illustrated in **a** (15 total groups, excluding the germline), colored according to **a** and grouped by order. Bottom, the largest fourth-order coefficients observed in the epistatic interaction model, with sites indicated. In both, error bars indicate standard error. **c**, CR-6261 force-directed graph, colored by a different set of 4 sites with the fewest strong linear effects and epistatic interactions (65, 66, 69, and 112.1). **d**, Higher-order significant epistatic contributions of CR-6261 mutation pairs, as in Fig. 3g, for binding H9. **e**, Scatterplot of significant epistatic interaction model coefficients for binding to H1 and H9. Terms at different orders are colored and sized as indicated. Selected coefficients are annotated. Significance in **d**, **e** indicates Bonferroni-corrected *p*-value < 0.05, see SI.

**ED Figure 8:**
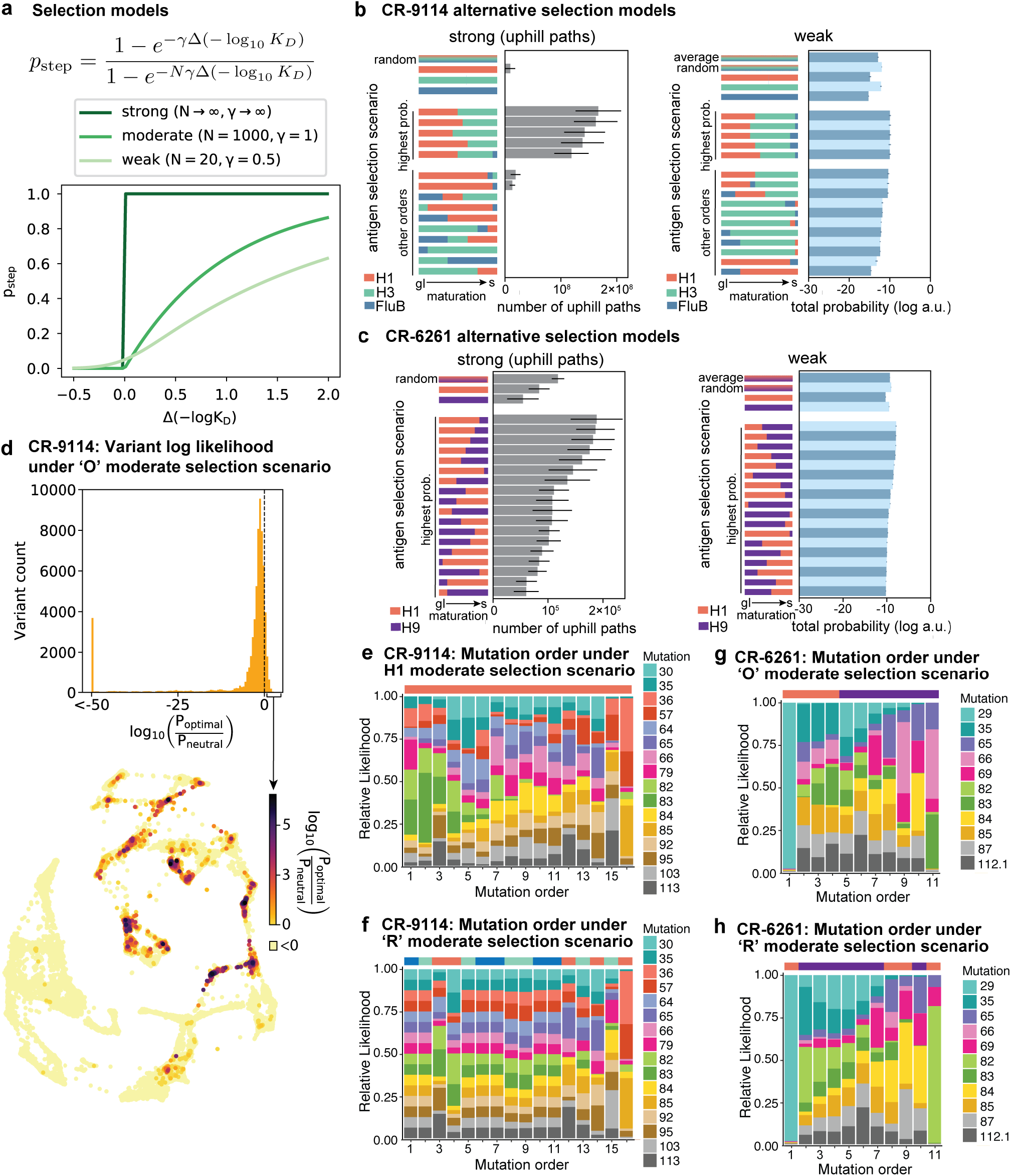
Likelihood of antigen selection scenarios and corresponding mutational pathways. **a**, Functional form of mutation step probability, illustrated for parameters chosen to represent strong, moderate, and weak selection models. **b, c**, Total log probability of the mutational trajectories between germline and somatic sequences for (**b**) CR-9114 and **(c)** CR-6261 under different antigen selection scenarios, assuming strong (left) or weak (right) selection, as shown for moderate selection in Fig. 4e,f. Strong selection scenarios are shown on a linear scale, as total probability is equal to the number of uphill paths. The “average” mixed scenario is not evaluated for strong selection, as the quantitative effect of averaging is undone by the binarizing effect of the transition probability. **ED Figure 8, continued: d**, Total log probability of each variant in the optimal sequential selection scenario. Top, histogram of the total probability of all pathways passing through each variant in the optimal selection scenario, divided by the total probability in a model with no selection, transformed to log10 scale. Dotted line indicates the 11% of variants favored in the selective model (log probability ratio greater than zero). Bottom, these favored variants are shown on the force-directed graph for CR-9114 H1 −logKd, as in Fig. 1g, with darker color according to the log probability ratio. Other variants with log probability ratio less than zero are shown in light yellow. **e–h**, Probability of mutation order assuming moderate selection, under antigen selection scenarios ‘H1’ (**e**) and ‘R’ (**f**) for CR-9114 and ‘O’ (**g**) and ‘R’ (**h**) for CR-6261, as in Fig. 4i,j.

