## Supplemental Information for "Binding affinity landscapes constrain the evolution of broadly neutralizing anti-influenza antibodies"

### Supplementary Information

**References** **29**

### 1 Antibody library production

#### 1.1 Yeast display plasmid and strains

To generate clonal yeast display strains and libraries for CR-9114, we cloned scFv constructs ($V_L$ -Ser(Gly$_4$Ser)$_5$-$V_H$-Myc) into the pCT302 plasmid [1] (kind gift from Dane Wittrup; Addgene #41845). For the clonal germline and somatic strains, gene blocks corresponding to the CR-9114 germline [2] or somatic [3] scFv sequence (codon optimized for expression in yeast) were cloned into pCT302 by Gibson Assembly [4] (plasmid maps in SI Files 4-5). For producing the plasmid backbone required for Golden Gate library generation (described below), we removed an existing Bsa-I site from the pCT302 plasmid by site-directed mutagenesis (Agilent #200521) and replaced the $V_H$ domain with the *ccdB* gene. To generate clonal yeast strains, Gibson Assembly products were transformed into electrocompetent DH10B *E. coli* cells, and the resulting plasmids were mini-prepped and Sanger sequenced. Following sequence confirmation, plasmids were transformed into EBY100 yeast cells as described in the high efficiency yeast transformation protocol [5]. Transformants were plated on SDCAA-agar and incubated at 30°C for 48 h, single colonies were restruck on SDCAA-agar and again incubated at 30°C for 48 h, and the resulting clonal yeast strains were verified to have the construct of interest by colony PCR. Construction of the yeast libraries is described below. All yeast strains were grown to saturation in SDCAA at 30°C, supplemented with 5% glycerol, and stored at -80°C.

CR-6261 clonal yeast display strains and libraries were generated in an identical manner to that of CR-9114, except where noted below (see SI Files 6-7 for plasmid maps corresponding to the germline and somatic sequences).

#### 1.2 Mutation selection

CR-9114 contains a total of 18 amino acid substitutions between the somatic variant and the reconstructed germline sequence. However, a library of $2^{18} = 262,144$ variants would be costly and time-consuming to produce and assay via our methods. We therefore identified 2 mutations that were distant from antigen contacts in the crystal structure [3]: A25S and E51D. We measured binding affinities for somatic sequences with and without these two mutations, and found that these variants had comparable affinities for both H1 and H3 (SI Fig. S1). Although these mutations may have some small impact on binding, especially in combination with others, excluding them allowed for a simpler cloning strategy and a more manageable library size.

Similar to the CR-9114 library design, we reduced the number of mutations present in the CR-6261 library by excluding 3 mutations that were distant from antigen contacts in the crystal structure: 6QE, L50P, and V101M [6]. We validated the marginal contribution of these mutations to binding by measuring the binding affinities for the somatic sequence with and without these mutations reverted to the respective germline residue (SI Fig. S2).

#### 1.3 Golden Gate assembly

For CR-9114, due to the number of mutations required and their positions along the heavy chain coding sequence, we designed a library cloning strategy using Golden Gate combinatorial assembly [7]. We divided the heavy chain coding region into 5 roughly equal fragments, ranging from 79 to 85 bp and each containing between 1 and 5 mutations. We added BsaI sites and additional overhangs to both ends of each fragment sequence, with cut sites carefully chosen so that the 5 fragments will assemble uniquely in their proper order within the plasmid backbone. For each fragment with $n$ mutations, we then ordered $2^n$ individual DNA duplexes with each possible combination of mutations (ranging from 2 to 32 versions for each fragment, a total of 66 fragments) from IDT (see SI File 2). By pooling the versions of each fragment in equal volumes, then pooling the 5 fragment pools in equimolar ratios, we obtained a randomized fragment mix containing all $2^{16}$ sequences present at approximately equal frequencies.

In addition to the fragment mix, we prepared the plasmid backbone for the Golden Gate reaction. We created a version of the yeast display plasmid with the counter-selection marker *ccdB* in place of the heavy chain sequence, with flanking BsaI sites (see above). We performed Golden Gate cloning using BsaI-HFv2 (NEB #R3733) following the manufacturer recommended protocol, with a 5:1 molar ratio of each fragment insert pool to plasmid backbone.

We transformed the assembly mix into electrocompetent *E. coli* (DH10B) via electroporation in 10 x 50 µL cell aliquots. We recovered each transformation in 5 mL SOC (2% tryptone, 0.5% yeast extract, 10 mM NaCl, 2.5 mM KCl, 10 mM $MgCl_2$, 10 mM $MgSO_4$, 20 mM glucose) at 37°C for 1h, and then transferred each to 100 mL of molten LB (1% tryptone, 0.5% yeast extract, 1% NaCl) containing 0.3% SeaPrep agarose (VWR #12001-922) spread into a thin layer in a 1L baffled flask (about 1 cm deep). The mixture was allowed to set on ice for an hour, after which it was kept for 18 hours at 37°C to allow for dispersed growth of colonies in 3D. We observed $\sim 3 \times 10^5$ colonies per aliquot, for a total of $\sim$ 3 million transformants. After mixing the flasks by shaking for 1h, we pelleted the cells and prepared plasmid by standard midiprep (ZymoPURE II Plasmid Midiprep, D4201), from which we obtained >120 µg of purified plasmid.

For CR-6261, we designed a library cloning strategy also using Golden Gate combinatorial assembly, but with fragments created by PCR instead of purchased. We divided the heavy chain coding region into 3 roughly equal fragments, each containing between 2 and 5 mutations. We designed these fragments such that the mutations they contain are close to the 3' or 5' ends and can thus be easily incorporated by PCR. PCR primers included mutations, BsaI sites, and unique overhangs chosen so that the 3 fragments would assemble uniquely in their proper order within the plasmid backbone. For each version of the three fragments, we generated dsDNA by PCR (52 PCR reactions in total; see SI File 3 for primer sequences). By pooling all versions of each fragment in equal volumes, then pooling the 3 fragment pools in equimolar ratios, we obtain a randomized fragment mix that, when ligated in the Golden Gate reaction, produces all of the $2^{11}$ sequences present at approximately equal frequencies.

In addition to the fragment mix, we prepared the plasmid backbone for the Golden Gate reaction. We created a version of the yeast display plasmid with the counter-selection marker ccdB in place of the 3-fragment sequence, with flanking BsaI sites. We performed Golden Gate cloning using BsaI-HFv2 (NEB #R3733) following the manufacturer recommended protocol, with a 7:1 molar ratio of fragment inserts to plasmid backbone.

The transformation of the CR-6261 library into *E. coli* was conducted in a similar fashion to that of CR-9114, except that 8x50 µL cell aliquots were transformed, and 600,000 colonies were pooled for plasmid midiprep.

#### 1.4 Yeast library production

We then transformed the CR-9114 plasmid library into EBY100 cells by standard high-efficiency protocols [5]. We recovered transformants in molten SDCAA (1.71 g/L YNB without amino acids and ammonium sulfate (Sigma-Aldrich #Y1251), 5 g/L ammonium sulfate (Sigma-Aldrich #A4418), 2% dextrose (VWR #90000-904), 5 g/L Bacto casamino acids (VWR #223050), 100 µg/L ampicillin (VWR # V0339)) containing 0.35% SeaPrep agarose (VWR #12001-922) spread into a thin layer (about 1 cm deep). The mixture was allowed to set on ice for an hour, after which it was kept for 48 hours at 30°C to allow for dispersed growth of colonies in 3D. From 5 such flasks, we obtained ~700,000 colonies (>10 times the library diversity). After mixing the flasks thoroughly by shaking for 1h, we grew cells in 5-mL tubes of liquid SDCAA for 5 generations and froze the saturated culture in 1-mL aliquots with 5% glycerol.

The CR-6261 yeast library was generated in a manner identical to that of CR-9114, except that ~60,000 colonies were pooled due to the smaller library size.

#### 1.5 Isogenic strain production

In addition to the full library, for both CR-9114 and CR-6261 we assayed a small number of variants by low-throughput flow cytometry for Tite-Seq validation. Any individual variant in the library can be produced in the same manner as described above: we simply selected the DNA duplex fragments corresponding to each desired variant and set up an individual Golden Gate reaction. The resulting assembled plasmid was transformed into *E. coli*, mini-prepped, and transformed into EBY100 in the same manner as described above. We verified the sequence identity of each variant by Sanger sequencing the entire scFv sequence.

We also constructed several strains for validation experiments that are not present in the full library. First, to test the impact of excluding mutations A24S and E46D from the CR-9114 library, we constructed a strain containing the remaining 16 somatic mutations by cloning a gene block of the corresponding $V_H$ sequence into the germline CR-9114 pCT302 plasmid via Gibson Assembly (SI Fig. S1). Second, to test the impact of the light chain on antigen binding, we constructed four strains comprising all possible combinations of the germline and somatic versions of the light and heavy chains by Gibson Assembly (SI Fig. S1). We note that all antibody libraries were constructed using somatic forms of the light chain, as these antibodies were isolated by combinatorial phage display [3, 8], and so it is not possible to infer the naturally paired germline light chain sequence.

For CR-6261, the revertant strains in the validation experiments described above (Section 1.2) were constructed by Gibson Assembly, as described for CR-9114. The CR-6261 light chain was previously determined not to impact binding [9].

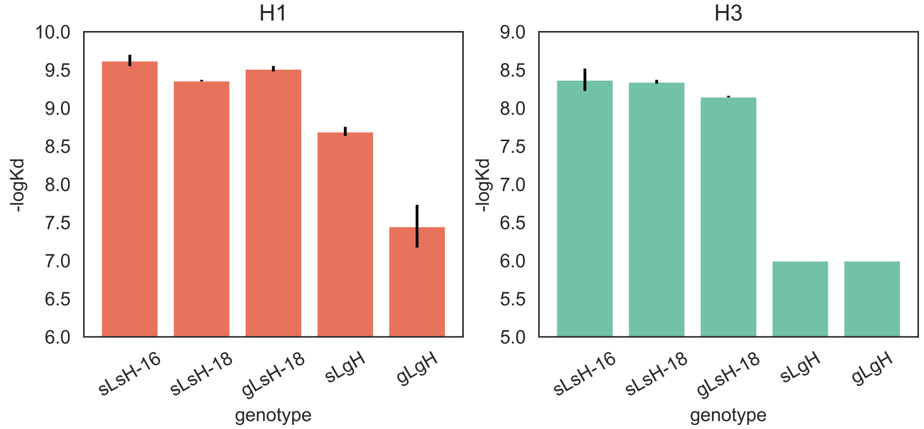

**Figure S1:** $-\log_{10} K_D$ for various combinations of CR-9114 somatic (s) and germline (g) light (L) and heavy (H) chains. Reversion of A24S and E46D (sLsH-16) compared to the fully mutated somatic sequence (sLsH-18) does not substantially impact binding affinity of CR-9114 to H1 (orange) or H3 (turquoise); these mutations are thus excluded from the CR-9114 library, as discussed in Section 1.2. Identity of the light chain does not impact H1 binding affinity when the heavy chain is somatic (gLsH-18 vs sLsH-18), but does when the heavy chain is germline (gLgH vs sLgH); in neither case is H3 binding affinity affected. Measurements made in biological duplicate; mean +/- SEM shown.

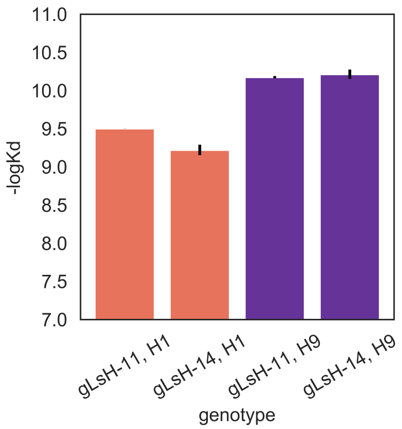

**Figure S2:** $-\log_{10} K_D$ for CR-6261 somatic reversions. Reversion of Q6E, L50P, and V101M (gLsH-11) does not substantially impact binding affinity compared to the fully somatic version of CR-6261 (gLsH-14) to H1 (orange) or H9 (purple); these mutations are thus excluded from the CR-6261 library, as discussed in Section 1.2. All variants have germline light chains (gL), as light chain identity does not impact affinity for CR-6261[9]. Measurements made in biological duplicate; mean +/- SEM shown.

### 2 Antigen production

#### 2.1 Choice of HA antigens

Both CR-6261 and CR-9114 were isolated from pooled PBMC from three donors who had received the trivalent 2006 influenza vaccine [8], which contained A/New Caledonia/20/1999 (H1N1), A/Wisconsin/67/2005 (H3N2), and B/Malaysia/2506/2004 (Victoria lineage) [10]. Because PBMC were isolated only 7 days after vaccination, it is unlikely that CR-6261 and CR-9114 matured in response to these specific antigens, and more likely that the vaccine elicited memory recall of these antibodies [11]. Here, we chose to measure binding affinities to diverse antigens spanning the range of breadth for both CR-9114 and CR-6261 (See Fig. S3 for HA subtype diversity). CR-9114 neutralizes strains across influenza A (groups 1 and 2) and influenza B, so we measured affinities to one strain from each of these groups, and selected vaccine-like strains: A/New Caledonia/20/1999 (H1N1), A/Wisconsin/67/2005 (H3N2), and B/Ohio/1/2005 (Victoria lineage). CR-6261 neutralizes strains across influenza A group 1, thus we measured affinities to two strains from distinct subtypes within group 1: A/New Caledonia/20/1999 (H1N1) and A/Hong Kong/1073/1999 (H9N2). We note that CR-9114 indeed binds A/Hong Kong/1073/1999 (H9N2) [3], but CR-9114 variant affinities for this strain were not measured here, as we prioritized measurements to antigens that span the breadth of each antibody.

#### 2.2 HA cloning, expression, and purification.

Hemagglutinin (HA) antigen was produced as previously described [3, 10, 13]. The HA ectodomain (Influenza A: residues 11–329 of HA1 and 1–176 of HA2 (H3 numbering); Influenza B: residues 1-523) of Influenza A/New Caledonia/1999 H1, Influenza A/Hong Kong/1999 H9, Influenza A/Wisconsin/2005 H3, and Influenza B/Ohio/2005, with N-terminal gp67 signal peptide and C-terminal biotinylation site (GGGLNDIFEAQKIEWHE), thrombin cleavage site, trimerization domain and His6 tag, were cloned into pFastbac (plasmid maps in SI Files 8-10). Recombinant bacmid was generated using the ThermoFisher Bac-to-Bac kit (ThermoFisher #10359016). Sf9 cells (ThermoFisher #B82501) were then transfected (ThermoFisher #A38915) with the resulting bacmids, and P0 HA-baculovirus was harvested 7 days post-transfection by clarifying viral supernatant at 1,000 x g for 10 min. HA-baculovirus was then amplified twice by successively infecting 187 million Sf9 cells with 100 µL of viral supernatant and incubating in a humidified incubator at 28°C for 12 days. To induce HA expression, 105 million High-Five cells (ThermoFisher #B85502) were resuspended with 15 mL P2 HA-baculovirus, incubated for 20 minutes at room temperature, and then transferred to a 1 L non-baffled flask with 200 mL Corning Express-Five media (ThermoFisher #10486025) supplemented with 200 mM L-glutamine (VWR #45000-676). Expression cultures were incubated in a shaking incubator at 28°C and 110 rpm for 48 hours, after which HA-containing media was clarified by spinning first at 1,000 x g for 5 min at 4°C, and then by spinning the resulting supernatant again at 4,000 x g for 20 min at 4°C. The clarified media was then dialyzed into PBS (VWR #45000-448) by performing 4 x 2-hour 10-fold buffer exchanges to remove metal chelators from culture media. Dialyzed media was then combined with 10 mL equilibrated NiNTA resin (Thermo #R90101), gently shaken for 3 hours at 4°C, and loaded onto a column. The resin was washed first with 15 column volumes Wash Buffer 1 (50 mM Tris pH 8 at 4°C, 300 mM KCl, 10 mM imidazole) and subsequently with 15 column volumes Wash Buffer 2 (50 mM Tris pH 8 at 4°C, 300 mM KCl, 20 mM imidazole). HA was eluted from the resin after 10 minutes incubation with Elution Buffer (50 mM Tris pH 8 at 4°C, 300 mM KCl, 250 mM imidazole). HA was then buffer exchanged into PBS using 10 kDa Amicon Ultra Centrifugal Filters (Millipore Sigma #UFC901008) and concentrated

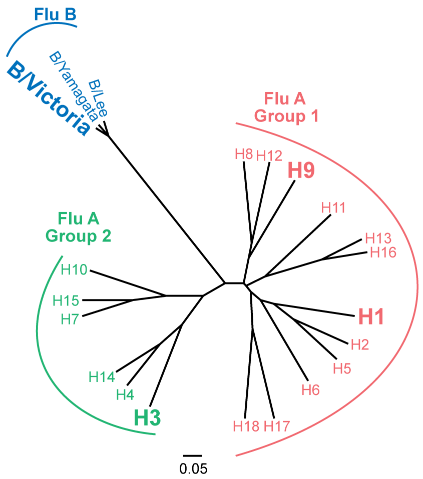

**Figure S3:** Hemagglutinin phylogenetic tree, adapted from [12]. Antigen subtypes used here (H1, H9, H3, and B/Victoria) shown in bold.

to at least 1 mg/mL for downstream biotinylation.

#### 2.3 BirA expression and purification.

BirA was expressed and purified as previously described [10]. Briefly, pET21a-BirA expression plasmid [14] (kind gift from Alice Ting; Addgene #20857) was transformed into BL21 (DE3). Transformed BL21 cells were grown in 4 L baffled flasks with 1 L low-salt LB medium (5 g/L NaCl, 5 g/L yeast extract (VWR #90000-722), 10 g/L tryptone (VWR #90000-286)) at 37°C to an OD (600 nm) of ∼0.8. The culture was then moved into cold water to bring it to 23°C, IPTG was added to a final concentration of 1 mM, and the culture was incubated at 23°C for ∼16 hours. The culture was then harvested by centrifugation (3,000 x g, 10 min), resuspended in 30 mL lysis buffer (50 mM Tris pH 8 at 4°C, 300 mM KCl, 10 mM imidazole, EDTA-free protease inhibitor cocktail tablet (Millipore Sigma #4693159001)), lysed by sonication (Branson Sonifier 450), and shaken at 4°C for 30 min. Lysate was clarified by spinning at 25,000 x g for 1h, and then the supernatant was incubated with 5 mL NiNTA resin at 4°C for 3 h with gentle shaking. The resin was pelleted by spinning at 500 x g for 5 min and washed twice by gentle shaking with 35 mL lysis buffer at 4°C for 30 min. Protein was eluted with 20 mL Elution Buffer (50 mM Tris pH 8 at 4°C, 300 mM KCl, 250 mM imidazole), buffer exchanged into Storage Buffer (50 mM Tris pH 7.5 at 4°C, 200 mM KCl, 5% glycerol) using 10 kDa Amicon Ultra Centrifugal Filters (Millipore Sigma #UFC901008), flash frozen in liquid nitrogen, and stored in single-use aliquots at -80°C.

#### 2.4 Biotinylation and HA-biotin quality control.

Purified hemagglutinin was biotinylated as previously described [10, 15]. Briefly, 100 µL HA (> 1 mg/mL) was incubated with 0.5 µL 1 M $MgCl_2$, 2 µL 100 mM ATP, 0.5 µL 50 mM biotin, and 2.5 µL BirA (10 mg/mL). This was mixed by gentle pipetting and incubated at 30°C with gentle rocking. After 1 h incubation, equivalent amounts of ATP, BirA, and biotin were added to the reaction, which was incubated for an additional hour at 30°C. Following the 2 h incubation, the 100 µL reaction was exchanged thrice into 15 mL PBS using a 50 kDa MWCO buffer exchange column (Millipore Sigma #UFC905008). The degree of biotinylation was then assessed by a streptavidin gel-shift assay, as previously described [15]. Briefly, 10-fold molar excess streptavidin (Millipore Sigma #189730) was added to 4 µg biotinylated HA and incubated at room temperature for 5 minutes prior to running on SDS-PAGE. Gels were transferred to nitrocellulose membranes and probed with mouse anti-His monoclonal antibodies (ThermoFisher #R930-25) and Goat-anti-mouse secondary antibodies (LiCor Cat#925-32210). HA was verified to be > 80% biotinylated by densitometry.

### 3 Tite-Seq assays

Tite-Seq was performed essentially as previously described [16], with some modifications as detailed below. For each antibody-antigen pair, three replicate Tite-Seq assays were performed on different days.

#### 3.1 Induction of antibody expression

On day 1, yeast scFv libraries, as well as germline and somatic clonal strains, were thawed by inoculating 5 mL SDCAA (1.71 g/L YNB without amino acids and ammonium sulfate (Sigma-Aldrich #Y1251), 5 g/L ammonium sulfate (Sigma-Aldrich #A4418), 2% dextrose (VWR #90000-904), 5 g/L Bacto casamino acids (VWR #223050), 100 µg/L ampicillin (VWR # V0339)) with 150 µL glycerol stock (saturated culture with 5% glycerol) and rotated at 30°C for 20 h. On day 2, yeast cultures were back-diluted to OD600 = 0.2 in 5 mL SDCAA and rotated at 30°C for approximately 4 h, or until reaching log phase (OD600 = 0.4 - 0.8). 1.5 mL log-phase cells were then pelleted, resuspended in 4 mL SGDCAA (1.71 g/L YNB without amino acids and ammonium sulfate (Sigma-Aldrich #Y1251), 5 g/L ammonium sulfate (Sigma-Aldrich #A4418), 0.2% dextrose (VWR #90000-904), 1.8% galactose (Sigma-Aldrich #G0625), 5 g/L Bacto casamino acids (VWR #223050), 100 µg/L ampicillin (VWR #V0339)), and rotated at room temperature for 20-22 h.

#### 3.2 Primary antigen labeling

On day 2, 20-22 hours post-induction, yeast cultures were pelleted, washed with 0.1% PBSA twice, and resuspended to an OD600 of 1. 700 µL of OD1 yeast cells were labeled with biotinylated HA at each of eleven antigen concentrations (half-log increments spanning 1 pM – 100 nM for H1 and H9, and 10 pM – 1 µM for H3 and influenza B, as well as no HA), with volumes adjusted such that the number of antigen molecules was in ten-fold excess of antibody molecules (assuming 50,000 scFv/cell). Yeast-HA mixtures were rocked at 4°C for 24 h.

#### 3.3 Secondary labeling

On day 3, yeast-HA complexes were pelleted by spinning at 3,000 x g for 10 minutes at 4°C, washed twice with 5% PBSA + 2 mM EDTA, and simultaneously labeled with Streptavidin-RPE (1:100, Thermo Fisher Cat#S866) and anti-cMyc-FITC (1:50, Miltenyi Biotec Cat#130-116-485) at 4°C for 45 minutes. Following secondary labeling, yeast were washed twice with 5% PBSA + 2 mM EDTA, and left on ice in the dark until sorting.

#### 3.4 Sorting and recovery

Yeast were sorted on a BD FACS Aria Illu, equipped with 405 nm, 440 nm, 488 nm, 561 nm, and 635 nm lasers, and an 85 micron fixed nozzle. Prior to sorting, single-color controls were used to compensate for the minimal FITC overlap with PE. Single cells were gated by FSC vs SSC, and then this population was sorted either by expression (FITC) or by expression and binding

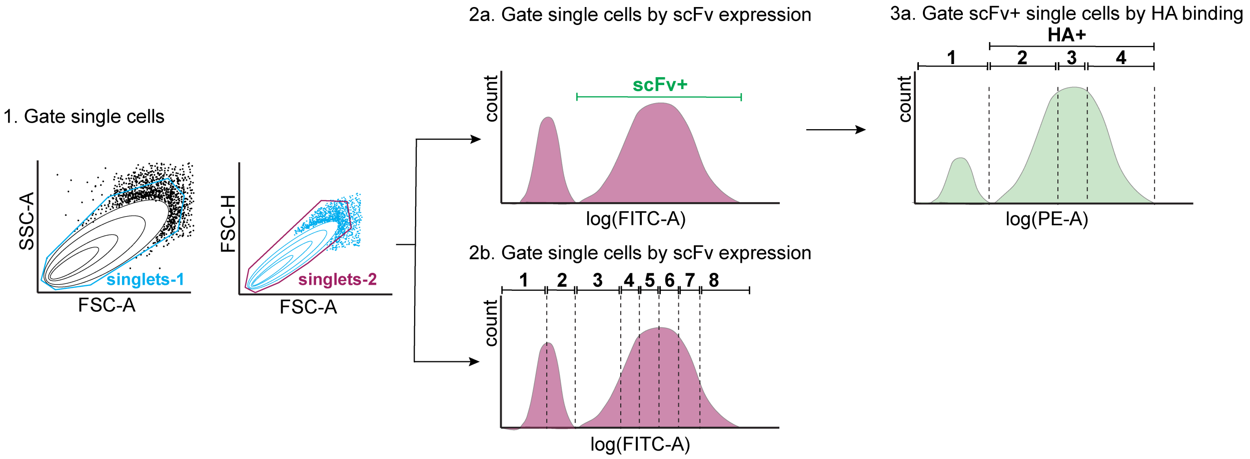

**Figure S4:** Tite-Seq sorting strategy. First, single yeast cells were gated by forward scatter (FSC) and side scatter (SSC) (step 1). Single cells were then either gated by scFv expression or HA binding. For the expression sort (step 2b), single cells were gated into eight bins along the log(FITC-A) axis, each containing 12.5% of the population. For the binding sort (steps 2a and 3a), scFv-expressing (scFv+) single cells were sorted into four bins along the log(PE-A) axis, with bin 1 comprising all HA- cells, and bins 2–4 each comprising 33% of the HA+ population.

(PE). For all sorts, at least ten-fold excess of the library diversity was sorted (~1.6 million cells for CR-9114; ~500,000 cells for CR-6261). For the expression sorts, singlets were sorted into 8 equivalent FITC log-spaced gates. For the binding sorts, FITC-positive cells were sorted into 4 PE bins (the PE-negative population comprised bin 1, and the PE-positive population was split into three equivalent log-spaced bins 2–4; see Fig. S4). Polypropylene collection tubes were coated and filled with 1 mL YPD supplemented with 1% BSA and placed on ice until recovery. Sorted cells were pelleted by spinning at 3,000 x g for 10 minutes, and supernatant was removed by pipette to avoid disturbing the pellets. Pellets were then resuspended in 4 mL SDCAA, a small amount was plated on SDCAA-agar to quantify recovery efficiency, and cultures were rocked at 30°C until reaching late-log phase (OD600 = 0.6 - 1.2).

#### 3.5 Sequencing library preparation

1.5 mL of late-log yeast cultures were pelleted and scFv plasmid was extracted using Zymo Yeast Plasmid Miniprep II (Zymo Research # D2004), per the manufacturer's instructions, and eluted in 10 µL elution buffer. Heavy-chain amplicon sequencing libraries were prepared by a two-step PCR as previously described [17]. In the first PCR, unique molecular identifiers (UMI), inline indices, and partial Illumina adapters were appended to the heavy chain through 3-5 amplification cycles to minimize PCR amplification bias. In the second PCR, the remainder of the Illumina adapter and sample-specific Illumina i5 and i7 indices were appended through 35 amplification cycles (see SI File 1 for primer sequences). The first PCR used 5 µL plasmid DNA as template in a 25 µL reaction volume, with Q5 polymerase according to the manufactuer's instructions (NEB # M0491L), and was incubated in a thermocycler with the following program: 1. 60s at 98°C, 2. 10s at 98°C, 3. 30s at 66°C, 4. 30s at 72°C, 5. GOTO 2, 2-4x, 6. 60s at 72°C. PCR products were then combined with carrier RNA and purified by 1.1X Aline beads (Aline Biosciences #C-1003-5), and eluted in 35 µL elution buffer. 33 µL of the elution was used as input for the second PCR, in a total volume of 50 µL using Kapa polymerase (Kapa Biosystems #KK2502) according to the manufacturer's instructions, and incubated in a thermocycler with the following program: 1. 30s at 98°C, 2. 20s at 98°C, 3. 30s at 62°C, 4. 30s at 72°C, 5. GOTO 2, 34x, 6. 300s at 72°C. The resulting sequencing libraries were purified by 0.85X Aline beads, amplicon size was verified to be ∼500 bp by running on a 1% agarose gel, and amplicon concentration was quantified by a fluorescent DNA-binding dye (Biotum #31068, per manufacturer's instructions). Amplicons were then pooled for each gate according to the number of sorted cells to ensure even sequencing coverage. The pool was further size-selected by a two-sided Aline bead cleanup (0.55-0.85X), and the final pool size was verified by Tapestation 5000 HS and 1000 HS. Final sequencing library concentration was determined by Qubit fluorometer and sequenced on an Illumina NovaSeq S2 (2x150) with 5% PhiX.

##### 3.6 Sequencing data processing

We first processed our raw sequencing reads to identify and extract the indexes and mutational sites, discarding priming regions and the constant regions between mutations. To do so, we developed custom Python scripts using the approximate regular expression library regex [18], which allowed us to handle complications in sequence parsing that arise from the irregular lengths of the indices and from sequencing errors. We accept sequences that match the entire read (with no restrictions on bases at mutational sites) within the following mismatch tolerances: 2 mismatches in the multiplexing index, 2 mismatches in the priming site, and 15 substitution mismatches within the 170 bases of constant antibody sequence.

We then examine the mutational sites to call germline or somatic alleles, producing binary genotypes ('0' for germline or '1' for somatic at each position). We require the exact germline or somatic sequence at every site: if there are any substitution errors in any of the mutation sites, the entire read is rejected. While it is possible to perform error correction based on Hamming distance to rescue reads with a few substitution errors, we find that on average only <8% of reads per sample contain any errors, and so we adopt the conservative approach of requiring perfect matching.

We next discarded sequencing reads with any mismatched indices (four total indices from the two PCR reactions), as well as reads with duplicate UMI sequences. Counts for each genotype were then tabulated, producing the final counts used for binding affinity inference (see Section 4). In Fig. S5 we show the mean and median coverage (reads per variant per concentration, after all filtering, averaged over all concentrations) for each antigen and replicate for both antibodies. On average we obtain a mean coverage of ∼350 for CR-9114 and ∼950 for CR-6261.

##### 3.7 Isogenic validation

Induction of scFv surface display, primary labeling, and secondary labeling of isogenic strains were performed identically to the Tite-Seq assay, except yeast cell and antigen volumes were scaled down by a factor of 10. Yeast cell FITC (scFv expression) and R-PE (HA binding) fluorescence intensity was assayed on a BD LSR Fortessa equipped with 4 lasers (440, 488, 561, and 633 nm). The equilibrium binding affinities ($K_D$) for each variant are inferred by fitting the log of a Hill function to the mean log R-PE fluorescence of scFv-expressing (FITC+) singlet yeast cells:

$$\text{mean log fluorescence} = \log_{10}\left(A_s \frac{c}{c + K_{D,s}} + B_s\right), \qquad (1)$$

where $c$ is the antigen concentration in molar units, $A_s$ is the increase in fluorescence due to saturation with antigen, $B_s$ is the background fluorescence, and $K_{D,s}$ is the equilibrium binding affinity. All isogenic measurements were performed in 2-3 biological replicates; see SI File 19 for isogenic $-\log_{10} K_D$.

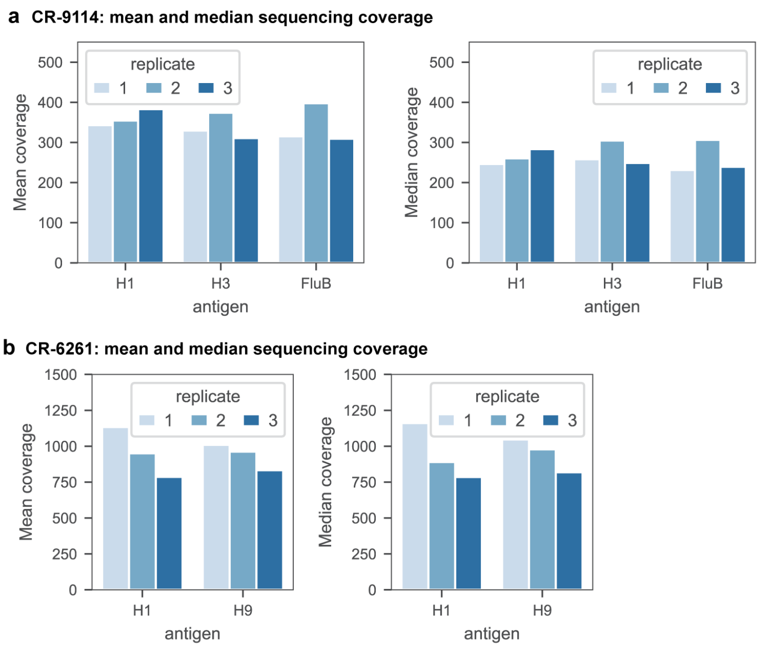

**Figure S5:** Sequencing coverage summary. Mean and median coverage (total reads per variant per concentration, after all filtering, averaged across all concentrations) for each antigen and each biological replicate, for **(a)** CR-9114 and **(b)** CR-6261.

### 4 Tite-Seq binding affinity inference

#### 4.1 Mean-bin approach

To infer binding affinities using a simple mean-bin approach [19], we incorporate sequencing data (the unique read counts of each genotype sequence $s$ in bin $b$ at concentration $c$, $R_{b,s,c}$) with flow cytometry data (the mean and standard deviation of $\log_{10}$-fluorescence of sorted cells in each bin $b$ at concentration $c$, $F_{b,c}$ and $\sigma_{F_{b,c}}$ respectively, and cell counts for each bin $b$ at each concentration $c$, $C_{b,c}$).

The mean fluorescence of each genotype sequence at each of the twelve antigen concentrations is calculated as:

$$\overline{F}_{s,c} = \sum_b F_{b,c}\, p_{b,s|c}, \tag{2}$$

where $p_{b,s|c}$ is the probability a cell with sequence $s$ would be sorted into bin $b$ at concentration $c$. $p_{b,s|c}$ is estimated from the sequencing read counts as:

$$p_{b,s|c} = \frac{\frac{R_{b,s,c}}{\sum_{s'} R_{b,s',c}} \cdot C_{b,c}}{\sum_{b'} \left( \frac{R_{b',s,c}}{\sum_{s'} R_{b',s',c}} \cdot C_{b',c} \right)}, \tag{3}$$

in other words, the fraction of total reads in the bin corresponding to sequence $s$, scaled by the number of sorted cells in that bin, normalized over the 4 bins for each concentration.

The uncertainty in the mean bin inference was propagated as:

$$\delta\overline{F}_{s,c} = \sqrt{\sum_b \left( \delta F_{b,c}^2\, p_{b,s|c}^2 + F_{b,c}^2\, \delta p_{b,s|c}^2 \right)}. \tag{4}$$

Here, $\delta F_{b,c}$ represents the spread in log-fluorescence values of cells sorted into the same bin $b$. While we could estimate this value using the bin width, in practice we find that the distribution of cell log-fluorescence values in a bin is far from uniform across the bin width. The distribution is often not normal either, but we find that approximating $\delta F_{b,c} \approx \sigma_{F_{b,c}}$, or the standard deviation in $\log_{10}$-fluorescence of cells sorted into bin $b$ at concentration $c$, adequately captures the typical variation. The error in $p_{b,s|c}$ arises largely from the sampling process of sequencing, which can be approximated as a Poisson process when read counts are relatively high. This gives

$$\delta p_{b,s|c} = \frac{p_{b,s|c}}{\sqrt{R_{b,s,c}}}. \tag{5}$$

Thus, $\delta \overline{F}_{s,c}$ can be written as

$$\delta \overline{F}_{s,c} = \sqrt{\sum_b \left( \sigma^2_{F_{b,c}}\, p^2_{b,s|c} + F^2_{b,c}\, \frac{p^2_{b,s|c}}{R_{b,s,c}} \right)}. \tag{6}$$

The equilibrium binding affinities ($K_D$) for each variant are inferred by fitting the logarithm of a Hill function to the resulting mean $\log_{10}$-fluorescence across the twelve antigen concentrations:

$$\overline{F}_{s,c} = \log_{10} \left( A_s \frac{c}{c + K_{D,s}} + B_s \right), \tag{7}$$

where $c$ is the antigen concentration in molar units, $A_s$ is the increase in fluorescence due to saturation with antigen, $B_s$ is the background fluorescence, and $K_{D,s}$ is the binding affinity. Fitting was performed with the *curve_fit* function of the Python package *scipy.optimize*. Reasonable bounds on the values of $A$ ($10^3$–$10^5$), $B$ ($10^0$–$10^3$), and $K_D$ ($10^{-14}$–$10^{-5}$) were imposed. Sequences leading to a failed optimization were deemed "non-binding".

Inferred $K_D$ outside of the titration boundaries were then pinned to the boundaries ($10^{-12}$ and $10^{-7}$ for H1 and H9; $10^{-11}$ and $10^{-6}$ for H3 and FluB). Inferred $K_D$ with high error (standard deviation of $\log_{10} K_D > 1.0$) or resulting from a poor fit ($r^2 < 0.8$) were removed from the data set prior to averaging $-\log_{10} K_D$ values across biological replicates.

#### 4.2 Maximum likelihood approach

In this approach we make the assumption that the fluorescence emitted by cells of a specific genotype is distributed log-normally, with parameters $\mu_{s,c}$ and $\sigma_{s,c}$ (the mean and standard deviation of the associated normal distribution respectively). At concentration $c$, a cell with genotype $s$ will fall into the bin $b$ ($\log_{10}$-fluorescence values $f_{s,c}$ ranging from $l_b$ to $h_b$) with probability:

$$P\left[l_b < f_{s,c} < h_b\right] = \int_{l_b}^{h_b} \frac{1}{\sqrt{2\pi\sigma^2_{s,c}}} e^{-\frac{1}{2}\left(\frac{f_{s,c}-\mu_{s,c}}{\sigma_{s,c}}\right)^2} \mathrm{d}f_{s,c} \tag{8}$$

$$= \frac{1}{2}\left( \mathrm{erf}\left(\frac{h_b - \mu_{s,c}}{\sigma_{s,c}\sqrt{2}}\right) - \mathrm{erf}\left(\frac{l_b - \mu_{s,c}}{\sigma_{s,c}\sqrt{2}}\right)\right). \tag{9}$$

Each cell sorted is an independent event, so the number of cells in each bin will be multinomially distributed, and thus the likelihood of sorting $n_{b,s|c}$ cells of sequence $s$ into bin $b$ at concentration $c$ is given by

$$\mathcal{L} = \prod_{s,c} \left(P\left[l_b < f_{s,c} < h_b\right]\right)^{n_{b,s|c}}, \tag{10}$$

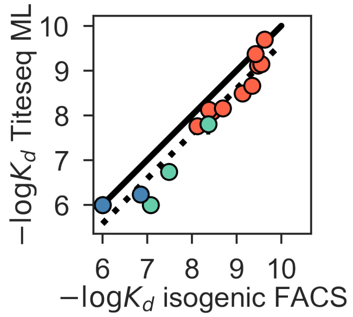

**Figure S6:** Correlation between $-\log_{10} K_D$ from ML inference on Tite-Seq data vs. $-\log_{10} K_D$ from isogenic flow cytometry. $-\log_{10} K_D$ to H1 (salmon), H3 (green), and Flu B (blue) shown for select variants, identical to those shown in ED Fig. 2d. Pearson's r = 0.97.

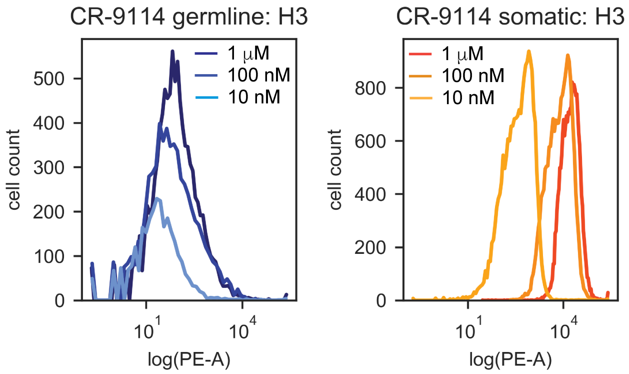

**Figure S7:** Distributions of PE-A fluorescence (HA binding) for isogenic CR-9114 strains incubated with H3. PE-A fluorescence distributions from flow cytometry of isogenic CR-9114 germline (left) and somatic (right) strains following incubation with 1 μM, 100 nM, and 10 nM H3, as described in Section 3.7. Shape of distribution varies for different clones and is not strictly log-normal, hence deviating from assumptions made in the maximum-likelihood binding affinity inference, as discussed in Section 4.2.

and the log-likelihood is

$$\log \mathcal{L} = \sum_{s,c,b} n_{b,s|c} \log P\left[l_b < f_{s,c} < h_b\right] \propto \sum_{s,c,b} p_{b,s|c} \log P\left[l_b < f_{s,c} < h_b\right]. \tag{11}$$

The probability $p_{b,s|c}$ is estimated as in the mean-bin approach (Section 4.1) and the log-likelihood is then maximized as a function of $\mu_{s,c}$ and $\sigma_{s,c}$ (BFGS method). The values of $A$, $K_D$, and $B$ are then estimated similarly as in Section 4.1, replacing $\overline{F}_{s,c}$ by $\mu_{s,c}$.

The $-\log_{10} K_D$ inferred by this ML approach correlate well with isogenic flow cytometry $-\log_{10} K_D$ (see Fig. S6), but not as well as those inferred by the mean-bin approach (ED Fig. 2d). The ML approach is predicated on the assumption that the fluorescence distribution for each variant is log-normal, which is often not the case (see Fig. S7). For these reasons, in addition to favoring a simple approach, we performed all subsequent analyses with $-\log_{10} K_D$ inferred by the mean-bin approach.

#### 4.3 Force-directed layouts

To represent the high-dimensional binding affinity landscape in two dimensions, we use a force-directed graph layout approach. Each sequence in the antibody library is a node, connected by edges to its single-mutation neighbors (sequences that can be reached by one additional somatic mutation). An edge between two sequences $s$ and $t$ is given the weight

$$w_{s,t} = \frac{1}{0.01 + \left|\log_{10}(K_{D,s}^{\text{ag}}) - \log_{10}(K_{D,t}^{\text{ag}})\right|}, \tag{12}$$

where $K_D^{\text{ag}}$ represent binding affinities to a particular antigen, ag. In the layouts shown in the main text, we use binding affinities to H1 for both CR-6261 and CR-9114. In force-directed layouts, edge weights correspond to the effective spring constant that tends to pull nodes closer together. Thus, a mutation from sequence $s$ to $t$ that has little impact on binding will cause that edge weight to be large, and the nodes will be pulled strongly together. A mutation from sequence $s$ to $t$ that causes a large difference in binding affinity (positive or negative) to the antigen will reduce the edge weight, moving those nodes further apart. After assigning all edge weights, we use the layout function *layout_drl* from the Python package *iGraph*, with default settings, to obtain the layout coordinates for each variant.

#### 4.4 Expression data

As noted above, antibody libraries were sorted into eight bins along the FITC-A fluorescence axis (where FITC-A fluorescence is proportional to expression), each comprising 12.5% of the total singlet population (Fig. S4). The expression mean bin was computed for each variant using the corresponding variant counts and fluorescence data, as described above for the mean-bin $K_D$ inference. These expression mean bin values were then averaged across all biological replicates for each antibody (9 replicates for CR-9114, 6 replicates for CR-6261), and correlation between biological replicates, as well as with $-\log_{10} K_D$ values, are illustrated in ED Fig 3. For the isogenic flow cytometry measurements, variant expression was computed as the mean log FITC-A fluorescence.

### 5 Epistasis analysis

#### 5.1 Linear interaction models

To infer specific mutational effects, we begin with simple linear models where the effects of mutations (and mutation combinations) add to produce phenotypes. Our log-transformed phenotypes for each variant $s$, $y_s = -\log_{10}(K_{D,s})$, are proportional to free-energy changes, and thus a natural null expectation is that they combine additively [20, 21] (although we also consider nonadditive epistatic interactions between individual loci here, and analyze the effects of an overall nonlinear transformation of this data in Section 5.4 below). Our additive-only model is

$$y_s = \beta_0 + \sum_{i=1}^{L} \beta_i x_{i,s} + \varepsilon, \tag{13}$$

where $L$ is the number of mutations for a given antibody, $\beta_0$ is an intercept term, $\beta_i$ is the effect of the mutation at site $i$, $x_{i,s}$ is the genotype of variant $s$ at site $i$, and $\varepsilon$ represents independently and identically distributed errors. Our general linear interaction models are

$$y_s = \beta_0 + \sum_i \beta_i x_{i,s} + \sum_{i<j}^{L} \beta_{ij} x_{i,s} x_{j,s} + \sum_{i<j<k}^{L} \beta_{ijk} x_{i,s} x_{j,s} x_{k,s} + \ldots + \varepsilon \quad (14)$$

where $\beta_{ij}$ represent second-order interaction coefficients between distinct sites $i$ and $j$, $\beta_{ijk}$ represent third-order interaction coefficients, and so on up to the desired maximum order of interaction.

There are multiple alternative coding systems for the binary genotypes $x_{i,s}$ that affect the values of inferred effects $\beta$ as well as their interpretation. Two common choices are (1) $x_{i,s} \in \{0, 1\}$, often called "biochemical" or "local" epistasis, and (2) $x_{i,s} \in \{-1, 1\}$, often called "statistical" or "ensemble" epistasis [22]. These frameworks are equivalent and related by a simple linear transformation, but the values of the coefficients vary between frameworks and have different interpretations. For ease of interpretation, in the main text and Extended Data figures we always show results obtained from inference in the biochemical epistasis framework. We discuss in Section 5.3 below the differences between these two frameworks, and present results from inference in the statistical epistasis framework.

For an antibody with $L$ mutations, there are $L$ possible orders of interactions, with a total of $2^L$ epistatic coefficients $\beta$. From a measurement of $y$ for all $2^L$ possible sequences, there is a simple linear transformation to calculate the resulting $2^L$ $\beta$ parameters [22]. This is a simple and fast approach to the calculation of epistasis that is widely used [23, 24]. However, we may instead wish to restrict our model to a lower order and examine whether it can explain the data with far fewer than $2^L$ parameters, as a conservative approach to detecting high-order epistasis.

Specifically, we truncate the model above at a maximum order $n$ and then fit and evaluate the resulting model. We begin with $n = 1$ and continue to increase $n$ until the optimal model has been identified. There are multiple strategies for selecting between models with different numbers of parameters, such as AIC and BIC; here we take a cross-validation approach. For each fold, we hold out 10% of the dataset, train models at each maximum order on the remaining 90%, and evaluate the prediction performance ($R^2$) of the model on the held-out test set. After averaging the performances across all 10 folds for each truncated model, we choose the order that maximizes the test set performance as the optimal maximal order of interaction. We then re-train the model truncated at this order on the full dataset to obtain the final coefficients. We find that the optimal model identified by cross-validation for each antibody-antigen pair always satisfies $p \ll N$, where $p$ is the number of model coefficients and $N$ the number of data points with measurable binding affinity (Fig. S8). This gives confidence that our parameter estimates are well constrained by the data, even in the absence of other regularization (such as Lasso or Ridge regularization approaches).

To train a model of given order on a set of sequences, we use ordinary least squares (OLS) regression with the Python package *statsmodels*. From this, we obtain the coefficient values $\beta$ with their standard errors and $p$-values. These coefficients and standard errors are used in Figures 2 and 3. To define significance of coefficients, we use a $p$-value cutoff of 0.05 with Bonferroni correction by the total number of model parameters. Coefficients, standard errors, $p$-values, and Bonferroni-corrected 95% confidence intervals are reported in Supplemental Files 14-18. We also predict phenotypes $\hat{y}$ for each sequence from the coefficients and use these values in Figure 4a,b.

#### 5.2 Structural analysis of epistatic coefficients

To examine the structural context of linear and pairwise coefficients, we performed two simple analyses. *(1)* We used PyMol[25] to count the number of HA residues within six angstroms of each antibody mutation site, and plot this number of proximal residues vs the linear effect of the

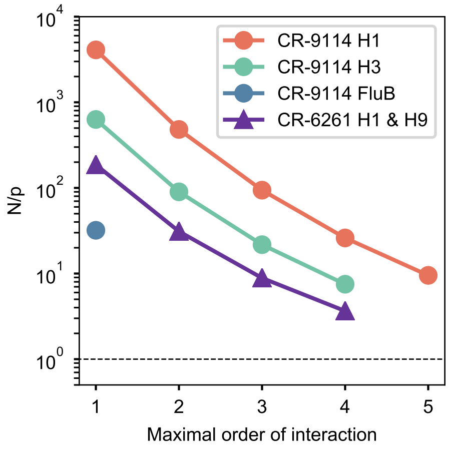

**Figure S8:** Scaling of number of datapoints $N$ compared to total parameter number $p$ for different antibody-antigen pairs at different maximal orders of interaction, up to the optimal maximal order for each antigen. Dotted line indicates $N/p = 1$, or equal numbers of parameters and datapoints.

corresponding mutation on HA binding (Fig. 2c; ED Fig. 5a). Six angstroms was chosen as an upper limit to capture potential antibody-antigen interactions[3, 6, 26–28], though we note that this analysis is robust to other distance thresholds. *(2)* We also used PyMol[25] to measure the distances between alpha carbons for all mutation pairs, and plotted these distances against the corresponding pairwise epistatic terms (Fig. 2f; ED Fig. 5b). We note that both of these analyses were performed with co-crystal structures of the somatic antibodies with HA (PDB ID: 4FQI (CR-9114–H5 (CR-9114–H1 crystal structure not available)[3]; 4FQY (CR-9114–H3)[3]; 3GBN (CR-6261–H1)[6]).

#### 5.3 Statistical epistasis and variance partitioning

The contrasting frameworks for epistasis are well described in [22]. In particular, a biochemical epistasis approach highlights one particular sequence as the "wildtype" or reference sequence and measures effects relative to its phenotype, whereas a statistical epistasis approach measures effects relative to the average background of all variants included. The biochemical approach benefits from easier interpretation of the coefficient magnitudes, particularly when there is a natural or relevant choice of reference sequence, but the coefficients at different orders are not statistically independent. The statistical approach allows for correct variance partitioning between interaction orders, but the interpretation of the coefficients can be sensitive to the set of sequences, particularly when not all possible sequences are represented or when a majority of sequences exhibit some uninteresting phenotype (e.g. lethal).

The results from statistical epistasis inference are shown in Fig. S9 for CR-9114 and Fig. S10 for CR-6261, in plots analogous to those in the main text and Extended Data. We find that the patterns of site participation in interactions are similar (although the coefficient magnitudes and signs are of course scaled differently). The group of five key sites discussed in Figure 3 (sites 30, 57, 65, 82, and 83 for CR-9114 binding to H1) exhibit coefficients that are significant for all 31 mutation combinations, consistent with the result from biochemical epistasis. Overall, the numbers of significant coefficients inferred in statistical epistasis models tends to be similar or slightly higher than for biochemical epistasis models, indicating that neither framework is a substantially more compact representation of epistasis than the other.

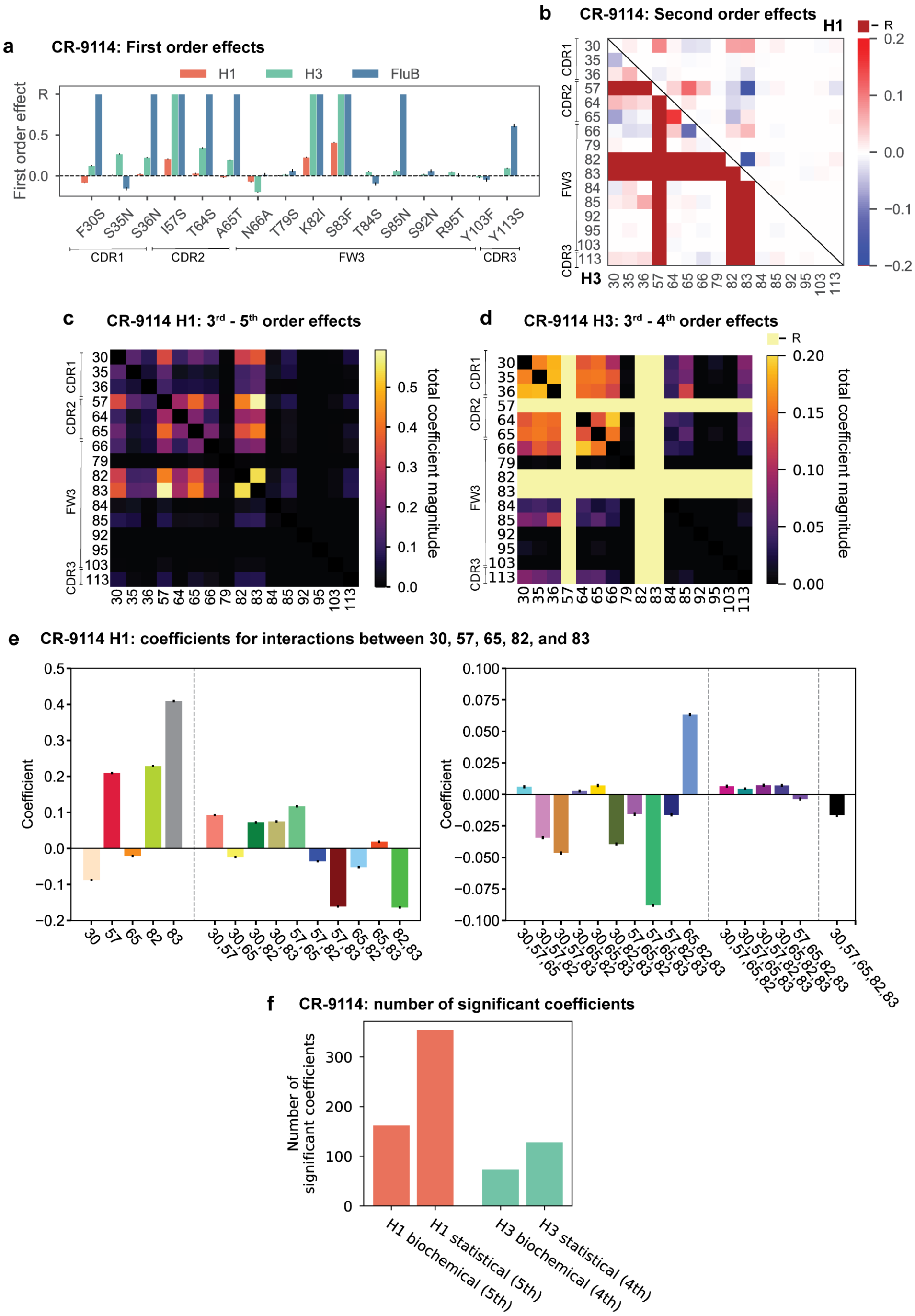

**Figure S9:** Results from statistical epistasis models for CR-9114. **a**, First-order effects, as in Figure 2a. 'R' indicates required mutations. **b**, Second-order effects for H1 (top right) and H3 (lower left), as in Figure 2d. Interactions with required mutations are noted in dark red. **c**, Cumulative higher-order effects for CR-9114 binding to H1, as in Figure 3a. **d**, Cumulative higher-order effects for CR-9114 binding to H3, as in Extended Data Figure 6f. **e**, Inferred interaction coefficients for the set of five key epistatic loci, as in Extended Data Figure 6b with corresponding colors. Note the different y-axis scales for the two subplots. Different interaction orders are separated by dotted lines. **f**, Number of significant coefficients at all orders for the biochemical and statistical epistasis models. The maximal order of interaction for each model is indicated in parentheses.

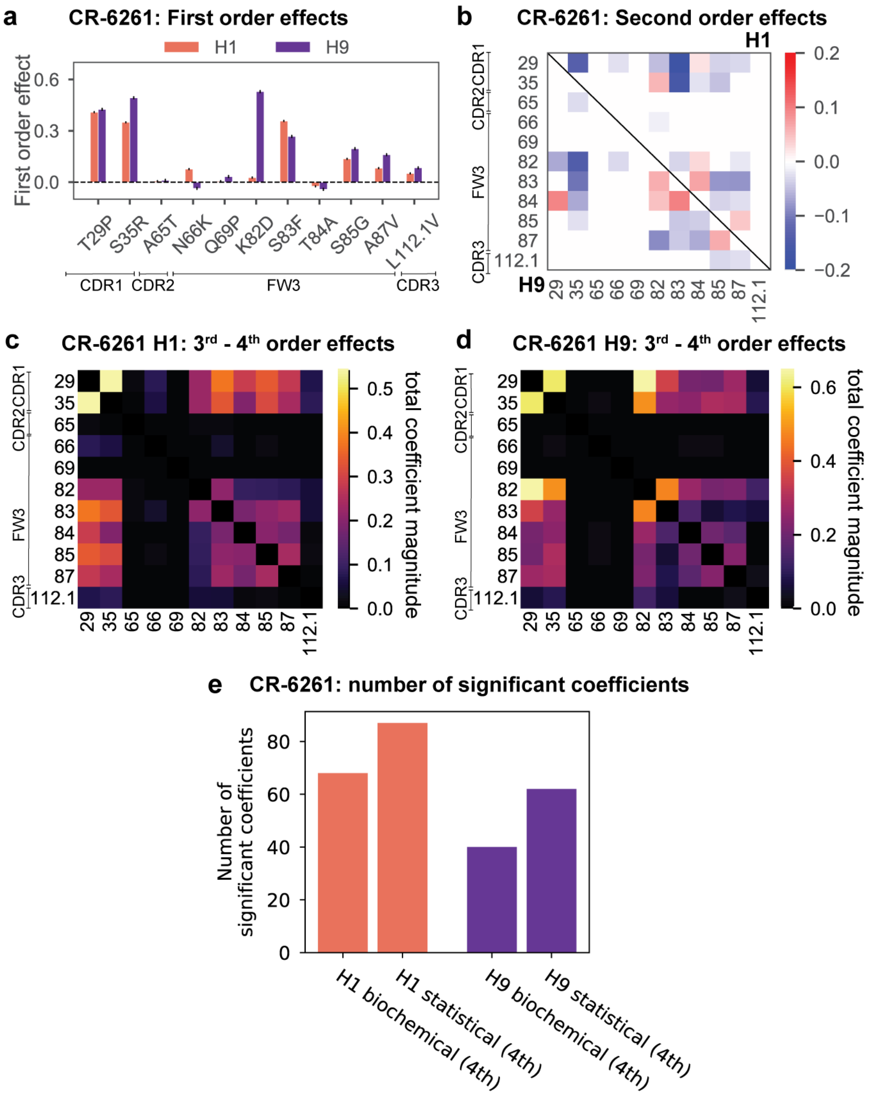

**Figure S10:** Results from statistical epistasis models for CR-6261. **a**, First-order effects, as in Figure 2b. **b**, Second-order effects for H1 (top right) and H9 (lower left), as in Figure 2e. **c**, Cumulative higher-order effects for CR-6261 binding to H1, as in Figure 3g. **d**, cumulative higher-order effects for CR-9114 binding to H9, as in Extended Data Figure 7d. **e**, number of significant coefficients at all orders for the biochemical and statistical epistasis models. The maximal order of interaction for each model is indicated in parentheses.

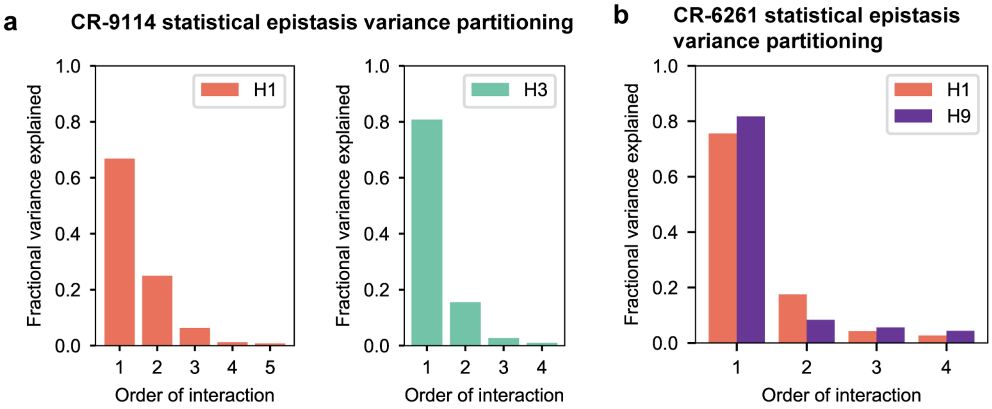

**Figure S11:** Variance partitioning of statistical epistasis models. **a**, Variance partitioning for CR-9114 binding to H1 (left) and H3 (right). **b**, Variance partitioning for CR-9114 binding to H1 and H9, denoted by colors as indicated.

In the statistical epistasis framework, we can also partition the variance explained by the model according to the interaction order. Here, we take the final inferred model at the optimal interaction order and evaluate the prediction performance ($R^2$) of each order as a fraction of the total performance of the full model. As shown in Fig. S11, we find that variance explained tends to decline with increasing order, as is also observed in some other protein epistasis datasets [23]. This indicates that interactions at higher order are more rare (compared to the total number of terms at each order, which scales combinatorially) and/or smaller in magnitude than those at lower order. However, this does not imply that rare, strong interactions of even higher order do not exist; for example, there may be some strong sixth-order interaction terms for CR-9114 binding to H1, but not enough to compensate for the many nonsignificant sixth-order terms in our cross-validation framework.

In particular, another alternative approach to the inference of epistasis is to infer a full $L^{\text{th}}$-order model rather than truncating to lower order. This approach calculates $2^L$ epistatic coefficients, one for every datapoint, which allows for the detection of strong interactions at any order with the caveat that many coefficients may simply reflect experimental noise, especially for higher-order terms. We follow the approach of [22, 24]: we calculate epistasic coefficients using a Walsh-Hadamard transform of the $-\log K_D$ values, and calculate standard errors on each coefficient via error propagation using the standard errors of the data. We define significant coefficients by a $p$-value cutoff of 0.05, with Bonferroni correction by the total number of parameters in the model (here $2^L$). We find that for all antibody-antigen combinations, this approach finds more significant coefficients than the optimal truncated models, many of which are at higher interaction orders than allowed in the truncated model (Fig S12). This analysis requires a measurement of $-\log K_D$ for every single variant, so we use data that has not been filtered for goodness-of-fit or error in the inference of binding affinity (see 4.1), including some sequences that have substantial error. Therefore we prefer to use the more conservative regression approach for our in-depth analysis of epistasis; this inference at full order confirms the existence, strength, and identity of the high-order interactions we discuss from the regression approach, while also indicating that additional and even higher-order terms may yet exist.

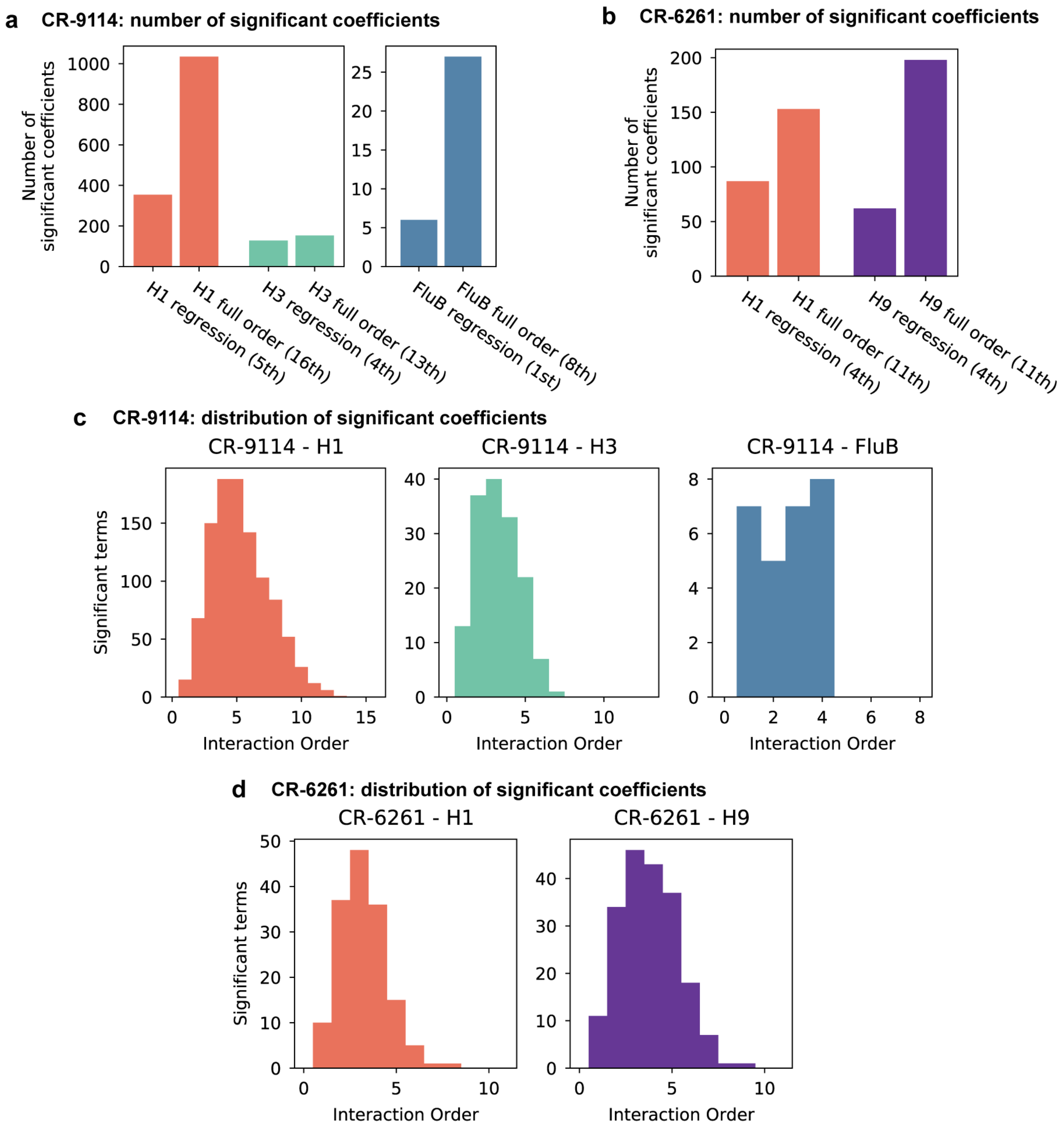

**Figure S12:** Epistasis inference at full order. **a**,**b**, Numbers of significant coefficients for the full-order inference compared to optimal truncated regression models for (**a**) CR-9114 and (**b**) CR-6261. Significance for both model types is determined by $p < 0.05$ with Bonferroni correction by the number of model parameters. **c**,**d**, Distribution of interaction orders of significant coefficients for (**c**) CR-9114 and (**d**) CR-6261.

#### 5.4 Nonlinear models

An alternative approach to understanding epistasis is to view nonlinearities in observed phenotype data as arising from a simple nonlinear transformation applied to an underlying, unobserved additive phenotype. In this view, a simple nonlinear "global epistasis" function with few parameters may describe the landscape as well or better than models of the sort described above, with their large number of "idiosyncratic epistasis" parameters. The underlying additive scale and the form of the nonlinear function can often be understood in a physical or biological sense: for example, mutations may contribute additively to protein stability, while protein stability is related to a measurable phenotype such as binding by a logistic-type function [29]. Many studies in other proteins have attempted to disentangle such global epistasis from idiosyncratic effects [29–34].

We already implement one global nonlinear transformation, by log-transforming our binding affinity measurements so that they are proportional to free energy changes, as described above. However, it is possible that another nonlinear transformation would capture the effects of many specific interaction coefficients, if there is a single underlying additive scale. In this section, we explore this possibility following the approach taken by [30]: we infer a nonlinear transformation that fits the phenotype data, invert it to "linearize" the phenotypes, re-fit interaction models on the linearized phenotypes, and then compare those model coefficients to the original coefficients to evaluate the role of the nonlinear transformation.

Our new model is

$$y_s = \Phi\left(y_{s,\text{add}}; k_m\right) = \Phi\left(\beta_0 + \sum_i^L \beta_i x_{i,s}; k_m\right), \tag{15}$$

where $y_s$ are the observed phenotypes ($-\log_{10} K_D$ values), $\Phi$ is a nonlinear function with a small number of associated parameters $k_m$, and $y_{s,\text{add}}$ are the underlying additive-scale phenotypes, parametrized as before by additive coefficients $\beta_i$.

To specify $\Phi$, we must choose a family of nonlinear functions. Typical choices include splines [29] or power transforms [23, 30]. We found that logistic (sigmoid) functions fit our data better than power transforms or splines, and they are monotonic and invertible. Specifically, our logistic function with four parameters is

$$\Phi(y; A, B, \mu, \sigma) = \frac{A}{1 + e^{\frac{(y-\mu)}{\sigma}}} + B. \tag{16}$$

Logistic functions capture two features that we observe: first, there is a saturation effect at low values of $-\log_{10} K_D$, corresponding to nonspecific binding that our measurements are unable to distinguish [35]; and second, for most antibody-antigen combinations we observe a saturation effect at moderately high values of $-\log_{10} K_D$. This latter effect is not due to limits on our measurement capabilities, as illustrated by higher values of $-\log_{10} K_D$ measured for the CR-6261 library to H9 compared to values of $-\log_{10} K_D$ measured for the CR-9114 library to H1, but instead due to widespread "diminishing returns" epistasis.

After specifying the functional form of $\Phi$, we must fit both the nonlinear parameters $k_m$ and underlying linear parameters $\beta_i$. In principle, one could fit all parameters jointly, using for example a maximum likelihood approach [29]. However, we take the simpler approach as implemented in the software package from [30], which first infers the additive parameters $\beta_i$ from the observed phenotypes and then infers the nonlinear function parameters $k_m$. We show the resulting fit of $\Phi$ in Fig. S13**a** for two representative examples, by plotting our estimate of the additive phenotypes $y_{s,\text{add}}$ on the x-axis and our observed phenotypes from data on the y-axis. We found that this simple procedure identified well-fitting $\Phi$ in a single step, and successive iterations did not significantly improve the fit.

After fitting the nonlinear transformation, we apply the inverse transformation to our observed phenotypes to obtain "linearized" phenotypes $y_{s,\text{lin}}$:

$$y_{s,\text{lin}} = \Phi^{-1}(y_s, k_m) \,. \tag{17}$$

Because the fit of $\Phi$ is not perfect, the linearized phenotypes $y_{s,\text{lin}}$ are not exactly equal to the estimated additive phenotypes $y_{s,\text{add}}$, although linear regression on both quantities produces extremely similar values of $\beta_i$. For values that lie above the domain of $\Phi^{-1}$, we pin them to the largest estimated additive phenotype.

Finally, we can take our linearized phenotypes $y_{s,\text{lin}}$ and infer interaction model coefficients $\beta'$ of various orders, exactly as described above for the untransformed "raw" phenotypes:

$$y_{s,\text{lin}} = \beta'_0 + \sum_i \beta'_i x_{i,s} + \sum_{i<j}^{L} \beta'_{ij} x_{i,s} x_{j,s} + \sum_{i<j<k}^{L} \beta'_{ijk} x_{i,s} x_{j,s} x_{k,s} + \ldots + \varepsilon. \tag{18}$$

We again perform this analysis in both the biochemical and statistical epistasis frameworks. If the inverse transformation has removed most or all of the nonlinearity, then the resulting optimal interaction models should be smaller (lower maximum order of interaction and/or fewer significant interaction coefficients).

Instead, we find that in all cases, the optimal order of interaction is unchanged or only decreased by one when inferring on linearized vs raw phenotypes. Specifically, the new (vs old) optimal orders are: 4th (vs 5th) for CR-9114 binding to H1, 4th (vs 4th) for CR-9114 binding to H3, 3rd (vs 4th) for CR-6261 binding to H1, 3rd (vs 4th) for CR-6261 binding to H9 in the biochemical epistasis framework, and 4th (vs 4th) for CR-6261 binding to H9 in the statistical epistasis framework. We can compare the numbers of significant coefficients in these optimal models inferred on linearized phenotypes to the models with the same maximum order inferred on raw phenotypes (Fig. S13**d**,**e**), where we see that the numbers are relatively comparable.

We next examine changes in the individual coefficients between these models. In Fig. S13**b**, we show two representative scatterplots between the raw phenotype coefficients $\beta$ and the linearized phenotype coefficients $\beta'$, where only significant coefficients are shown for clarity. While some coefficients show dramatic changes, overall the two sets of coefficients are quite well correlated. To see which sites are involved in strong changes, we can also represent coefficient changes in a heatmap format (Fig. S13**c**). Here, diagonal cells show the change in coefficient for single sites ($\beta'_i - \beta_i$), while off-diagonal cells show the sum of coefficient changes over all pairwise and higher terms involving each pair of mutations. We observe that for some antibody-antigen pairs, such as CR-9114 binding to H1, the strongest net changes are negative, though not negative enough to remove the many significant coefficients. For other antibody-antigen pairs such as CR-6261 binding to H1, there are both positive and negative net changes, indicating that the nonlinear transformation is changing the epistatic landscape rather than correcting for it.

In summary, we find that nonlinear logistic transformations can account for a portion of the nonlinearities observed in our data, sometimes reducing the maximal order of interaction by one. However, all antigen-antibody pairs still exhibit strong idiosyncratic epistasis up to at least 3rd order after correcting for global epistasis, and the resulting numbers and magnitudes of significant coefficients are not drastically changed. Thus, it does not appear that global epistasis can explain our data much more simply than models with individual interactions, and so we confine our main analysis to idiosyncratic epistasis models.

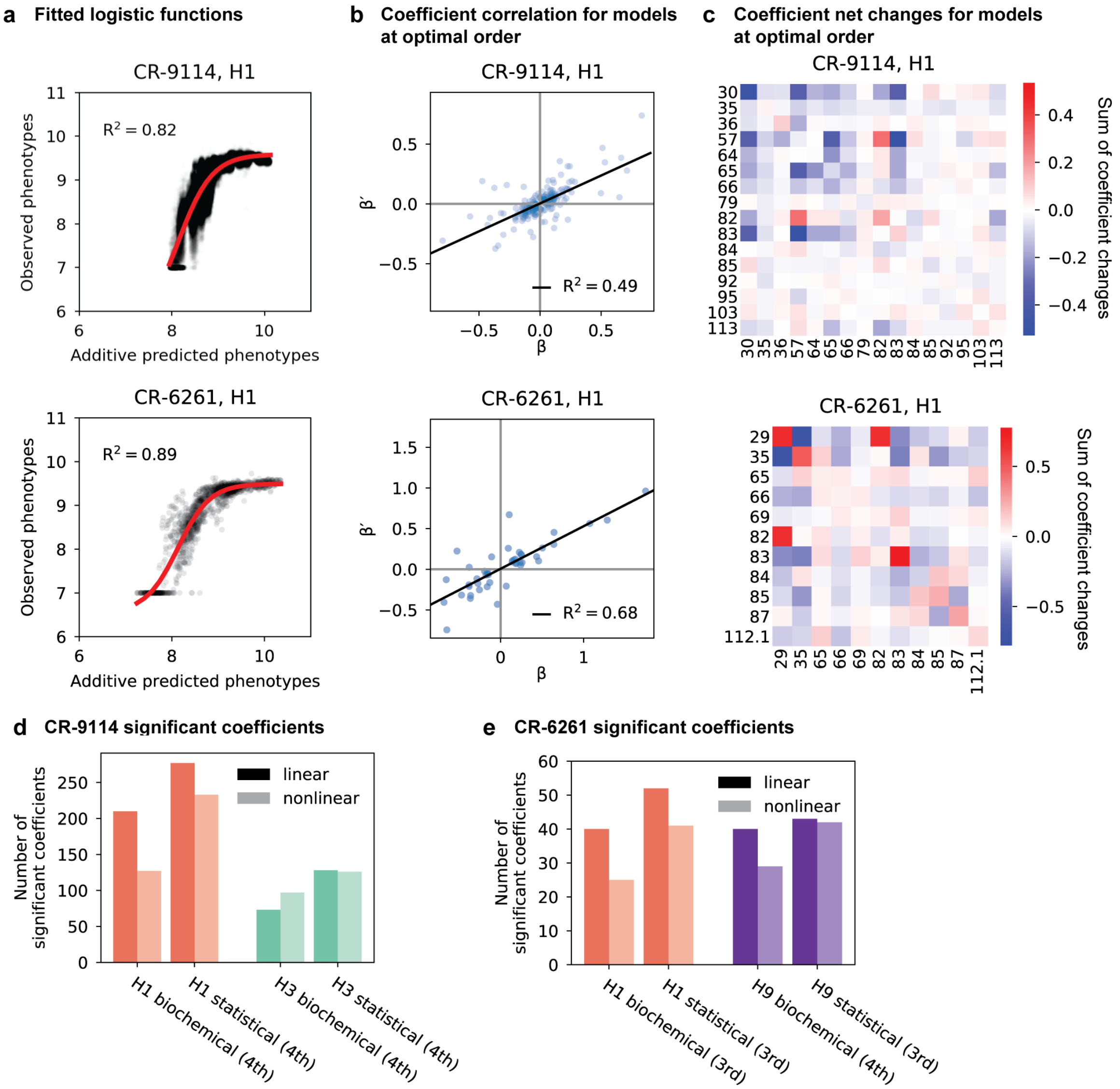

**Figure S13:** Results from epistasis models with nonlinear transformations. **a**, Fitting logistic functions to additive predicted phenotypes. Red lines indicate the optimized logistic function $\Phi$, with $R^2$ as indicated. **b**, Scatterplot of coefficients $\beta'$ from the optimal order model inferred on linearized data (after inverting the best-fit nonlinear transformation) against original coefficients $\beta$ for the model with the same maximum order. **c**, Net changes of coefficients by site. Diagonal cells show changes in linear coefficients. Off-diagonal cells show the sum of changes over terms at all orders (2nd and above) in which the given pair of mutations is involved. For **a**-**c**, we show two representative antibody-antigen combinations: CR-9114 binding to H1, top, and CR-6261 binding to H1, bottom. **d**-**e**, Number of significant coefficients in optimal order models fit to phenotypes transformed by the inverse nonlinear function (light bars), compared to original coefficients from linear models with the same maximal order (dark bars), for (**d**) CR-9114 and (**e**) CR-6261. The epistasis type and model order are indicated on the x-axis.

### 6 Pathway analysis

#### 6.1 Selection models

To study the likelihood of various mutational pathways leading from the germline to the somatic sequence, we must assume a selection model. Selection in germinal centers is considerably more complex than in classical population genetics models, involving spatial structure, changing population sizes, and T-cell mediated selection, among other factors [36]. Capturing these aspects in quantitative models is an active field of research [37]. However, here we wish to adopt an extremely simple model of selection as a first step in understanding the impacts of the binding affinity landscape on antibody selection, with the goal of understanding the implications of the expectation that mutational steps become more probable as their effect on binding affinity becomes more positive. Combining the more realistic models of immune selection with our detailed characterization of mutational effects on antigen binding affinity remains an interesting avenue for future work.

Here, we restrict to the weak-mutation regime where mutation fixation events occur independently of one another. Selection proceeds as a Markov process, where the population is characterized by a single sequence that acquires a single mutation at each discrete step [38]. We choose a simple form for the fixation probability of a mutation from sequence $s$ to sequence $t$, as discussed below. This then determines the transition probability for the population to move from $s$ to $t$. We assume that sequences cannot back-mutate (i.e. a residue changing from the somatic allele to the germline allele), and do not acquire multiple mutations in the same step. The absence of back-mutation is justified by the relatively large number of possible mutation sites compared to the total number of mutation events.

We define the transition probability of a single mutational step from the classical fixation probability for a mutation with selection coefficient $\sigma$ in a population of size $N$ [39]:

$$p_{\text{step}}(\sigma, N) = \frac{1 - e^{-\sigma}}{1 - e^{-N\sigma}}. \tag{19}$$

Here we define the selection coefficient $\sigma$ to be proportional to the difference in log binding affinities to a particular antigen between the two sequences $s$ and $t$:

$$\sigma = \gamma \Delta^{\text{ag}}_{s,t} = \gamma(-\log_{10} K^{\text{ag}}_{D,t} - (-\log_{10} K^{\text{ag}}_{D,s})). \tag{20}$$

This model has two tunable parameters: $N$ represents the effective population size and $\gamma$ represents how strongly differences in binding affinity impact fitness. We chose three parameter values to span a range of selection strengths (see ED Fig. 8): moderate, with $N = 1000$ and $\gamma = 1$; weak, with $N = 10$ and $\gamma = 0.3$; and strong, with $N \to \infty$ and $\gamma \to \infty$ such that $p_{\text{fix}}$ reduces to a step function (1 if $\Delta > 0$ and 0 otherwise). These three models all show similar results, with differences between selection scenarios becoming more exaggerated with stronger selection and less exaggerated with weaker selection, as expected (see ED Fig. 8).

From the fixation probabilities for a given parameter regime, we can compute the transition probability (up to a constant factor) for all sequences $s, t$ over all antigens ag,

$$P^{\text{ag}}_{s,t} = \begin{cases} p_{\text{step}}(\Delta^{\text{ag}}_{s,t}, \gamma, N) & \text{if } t \text{ has one more somatic mutation than } s \\ 0 & \text{otherwise} \end{cases},$$

which we use for all of the calculations described below for results presented in Figure 4 and Extended Data Figure 8.

#### 6.2 Scenario, mutation, and variant probabilities

It is particularly useful to store the probabilities $P^{\text{ag}}_{s,t}$ as (sparse) transition matrices $P^{\text{ag}}$ of dimension $2^N \times 2^N$ for each antigen, where entries are nonzero only where sequence $t$ has one more somatic mutation than $s$.

First, we wish to obtain a measure of total probability for a particular antigen scenario, as shown in Figure 4c,d. We calculate this by computing the matrix product over all mutational steps $i$ for a particular sequence of antigen contexts $\{\text{ag}_1, \dots, \text{ag}_L\}$:

$$\mathcal{P}_{tot} = \sum_{\text{paths}} \left( \prod_{\text{steps}} P_{\text{step}} \right) = \left[ \prod_{i=1}^{L} P^{\text{ag}_i} \right]_{s_{\text{g}}, s_{\text{s}}}, \tag{21}$$

where $[\cdot]_{s,s'}$ corresponds to taking the matrix element in the row corresponding to variant $s$ and column corresponding to variant $s'$. In the right-most term, the products are matrix operations and $s_{\text{g}}$, $s_{\text{s}}$ are respectively the indices of the germline and somatic variants.

We note that the transition probabilities $P^{\text{ag}_i}$ are not normalized at each step. In practice, this means that mutations are optional: many outcomes will not reach the somatic sequence and the likelihood encodes the probability of reaching the somatic state. This makes it possible to compare different scenarios, as some scenarios are more likely than others to reach the somatic state. However, because these values do not represent true probabilities — the units are arbitrary — they cannot be compared between antibodies or between selection models. The exception is for the strong scenario, where the total probability for each path is 1 if all steps are uphill ($\Delta^{\text{ag}}_{s,t} > 0$) and 0 otherwise. Thus, here $\mathcal{P}_{tot}$ has a natural interpretation as the total number of uphill paths. When we show results from the strong model (Figure 4c,d and Extended Data Figure 8b,c), we represent uphill path numbers on a linear scale without log-transforming.

Although there are many possible antigen exposure scenarios, we restrict our analysis to several classes. First, in single-antigen scenarios, all steps $i$ use the same antigen. Second, for sequential scenarios, antigen exposures must occur in non-repeating segments (for example, H1 - H3 - H1 is not allowed), although we consider all possible lengths and orders of segments.

Mixed scenarios are more complicated, as we do not fully understand the nature of B cell interactions with multiple antigens in the same germinal center [40–42]. One option is to assume that the B cell engages the antigen for which it has the highest affinity and define $\Delta$ by the maximum binding affinity across all possible antigens at each step, but this definition would trivially imply that the mixed scenario has the highest probability. Instead, we choose two alternatives: first, "average" mixed, where we assume the B cell engages all antigens and use the average binding affinity change over all three (for CR-9114) or two (for CR-6261) antigens, $\Delta_{\text{mixed}} = \frac{1}{N_{\text{ag}}} \sum_{\text{ag}} \Delta_{\text{ag}}$; and second, "random" mixed, where we assume the B cell randomly engages a single antigen and hence the antigen at each mutational step is chosen randomly. For the latter definition, we calculate $\mathcal{P}_{tot}$ as described above for 1000 randomly drawn scenarios and average the resulting log probability. When we illustrate mutational paths and mutation orders, we choose the scenario with median probability from the 1000 random draws.

We estimate the error of these probabilities by bootstrapping. Specifically, for 10 bootstrap iterations, we resample each binding affinity $-\log_{10} K^{\text{ag}}_{D,s}$ from a normal distribution according to its value and standard deviation. We then recalculate the total probability $\mathcal{P}_{tot}$, transform by the natural log, and average over the 10 values to obtain mean and s.e.m. values as shown in Figure 4 and ED Figure 8. We note that for the strong selection scenario (where probabilities represent total numbers of uphill paths), values are not log-transformed, and many scenarios have total path numbers of exactly zero. For CR-6261, all mutations at the first mutational step are neutral (with the exception of one mutation that improves affinity for H1 only), and so we allow all mutations with equal probability for the first step in the strong selection model.

Next, we wish to obtain the probability that a mutation at site $m$ happened at a specific step $j$ (Fig. 4g,h). As we are focusing on one antigen context, we can normalize the transition matrices and define:

$$\tilde{P}^{\text{ag}}_{s,t} = P^{\text{ag}}_{s,t} \times \left( \sum_t P^{\text{ag}}_{s,t} \right)^{-1}, \tag{22}$$

if $P^{\text{ag}}_{s,t} \neq 0$ and 0 otherwise. We can further restrict the transition matrix at step $j$, $\tilde{P}^{\text{ag}_j}$, to have nonzero probability only when the mutation that occurs is at a particular residue $\alpha$, $\tilde{P}^{\text{ag}_j}_{\alpha}$. The total relative probability for that site at that mutational step under an antigen exposure scenario is then

$$\mathcal{P}_{j,\alpha} = \left[ \left( \prod_{i=1}^{j-1} \tilde{P}^{\text{ag}_i} \right) \cdot \tilde{P}^{\text{ag}_j}_{\alpha} \cdot \left( \prod_{i=j+1}^{L} \tilde{P}^{\text{ag}_i} \right) \right]_{s_{\text{g}},s_{\text{s}}}. \tag{23}$$

Because a sequence of $L$ steps starting from the germline can only lead to the somatic state, $\tilde{P}$ verifies $\left[ \prod_{i=1}^{L} \tilde{P}^{\text{ag}_i} \right]_{s_{\text{g}},s_{\text{s}}} = 1$. With the relation $\sum_\alpha \tilde{P}^{\text{ag}_j}_{\alpha} = \tilde{P}^{\text{ag}_j}$ this implies that these probabilities are already normalized: $\sum_\alpha \mathcal{P}_{j,\alpha} = 1$.

We include these results for the "random" mixed scenario in the main text, as this scenario better captures the frustration resulting from competing selection forces to bind distinct antigens [40, 41]. We find that for CR-6261, mutation and pathway probabilities are quite similar for these two mixed scenarios, due to the correlated improvements in binding both antigens. For CR-9114, we find that the "random" mixed scenario is less likely to have led to the somatic sequence than the "average" mixed scenario. This is because in the "random" mixed scenario, the sequence often cannot improve in binding when faced with H3 or influenza B selection early on. The "average" mixed scenario favors paths that are also favored in the most likely sequential scenario, albeit less strongly, because the improvements in binding to one antigen are often diluted by averaging with other antigens. We refrain from studying the "average" mixed scenario for strong selection because it is essentially equivalent to choosing the antigen with maximum improvement: the quantitative effect of averaging is undone when the transition probability is binarized.

Next, we wish to determine the total probability of each variant (Extended Data Figure 8d), i.e. the sum of probabilities of all paths passing through that variant. For a variant $s$ that contains $j$ mutations, we calculate

$$\mathcal{P}_s = \left( \left[ \prod_{i=1}^{j} \tilde{P}^{\text{ag}_i} \right]_{s_{\text{g}},s} \right) \cdot \left( \left[ \prod_{i=j+1}^{L} \tilde{P}^{\text{ag}_i} \right]_{s,s_{\text{s}}} \right), \tag{24}$$

where the first term is the probability of reaching sequence $s$ at mutational step $j$, and the second term is the probability of reaching the somatic sequence after passing through sequence $s$. When representing this number we add an additional normalisation factor, $\mathcal{P}'_s = \mathcal{P}_s \times n_j$, where $n_j = \binom{L}{j}$ is the number of sequences with $j$ mutations, so that variants with different numbers of mutations have comparable values. $\mathcal{P}'_s$ thus represents the ratio of the probability in a selective model to the probability in a neutral model (which is $1/n_j$).

Finally, to identify the most likely paths under a given exposure scenario, we reframe this Markov process as a directed weighted graph. Each sequence $s$ is a node, and a directed edge exists towards all sequences $t$ that can be reached by one additional somatic mutation. The edge weight is calculated from the transition probability, $w_{s \to t} = -\log(P^{\text{ag}}_{s,t} + \epsilon)$, where $\epsilon$ is an extremely small value to ensure weights are finite. In this graph framework, we can use fast algorithms to obtain the "shortest" paths from the germline to the somatic node (those for which the sum of weights is lowest, i.e. the total probability is highest). Specifically, we use the *shortest_simple_paths* function from the Python package *networkx* [43] to compute the $k$ shortest paths, as shown in Figure 4e,f and Extended Data Figure 8. This method is exact and uses the algorithm described in [44].

[2] Veronique Giudicelli, Patrice Duroux, Chantal Ginestoux, Geraldine Folch, Joumana Jabado-Michaloud, Denys Chaume, and Marie-Paule Lefranc. IMGT/LIGM-DB, the IMGT® comprehensive database of immunoglobulin and T cell receptor nucleotide sequences. *Nucleic Acids Research*, 34(suppl_1):D781–D784, 2006.

[13] Irina Margine, Peter Palese, and Florian Krammer. Expression of functional recombinant hemagglutinin and neuraminidase proteins from the novel H7N9 influenza virus using the baculovirus expression system. *JoVE (Journal of Visualized Experiments)*, (81):e51112, 2013.

[14] Mark Howarth, Keizo Takao, Yasunori Hayashi, and Alice Y Ting. Targeting quantum dots to surface proteins in living cells with biotin ligase. *Proceedings of the National Academy of Sciences*, 102(21):7583–7588, 2005.

[15] Michael Fairhead and Mark Howarth. Site-specific biotinylation of purified proteins using BirA. In *Site-Specific Protein Labeling*, pages 171–184. Springer, 2015.

[16] Rhys M Adams, Thierry Mora, Aleksandra M Walczak, and Justin B Kinney. Measuring the sequence-affinity landscape of antibodies with massively parallel titration curves. *eLife*, 5:e23156, dec 2016.

[17] Alex N Nguyen Ba, Ivana Cvijović, José I Rojas Echenique, Katherine R Lawrence, Artur Rego-Costa, Xianan Liu, Sasha F Levy, and Michael M Desai. High-resolution lineage tracking reveals travelling wave of adaptation in laboratory yeast. *Nature*, 575(7783):494–499, 2019.

[18] Matthew Barnett. regex.

[25] Schrödinger, LLC. The PyMOL molecular graphics system, version 1.8. November 2015.

[37] Assaf Amitai, Luka Mesin, Gabriel D. Victora, Mehran Kardar, and Arup K. Chakraborty. A population dynamics model for clonal diversity in a germinal center. *Frontiers in Microbiology*, 8, 2017.
